# Image-scanning light-sheet microscopy for high-speed volumetric imaging of complex biological dynamics

**DOI:** 10.64898/2026.04.07.716805

**Authors:** Yusuke Tomina, Ayumu Ishijima, Yu Toyoshima, Hikaru Shishido, Ryu Hirooka, Kazuki Mukumoto, Chentao Wen, Manami Kanamori, Koyo Kuze, Yuko Murakami, Suzu Oe, Sae Tanaka, Yusuke Yonamine, Yukinori Nishigami, Keisuke Goda, Kuniharu Ijiro, Toshiyuki Nakagaki, Kazuharu Arakawa, Takeshi Ishihara, Shuichi Onami, Yuichi Iino, Hideharu Mikami

## Abstract

Volumetric fluorescence microscopy is a powerful method for studying complex biological systems because it enables comprehensive observation of structural and physiological dynamics. In particular, light-sheet microscopy (LSM) is a leading option for real-time volumetric fluorescence imaging as it combines high imaging speed, low phototoxicity, minimal photobleaching, high spatiotemporal resolution, and low computational burden. To capture fast biological events, various efforts have been made to improve the imaging speed of volumetric fluorescence microscopy, including LSM. However, existing approaches entail significant trade-offs that make routine volumetric imaging at and beyond video rates challenging under practical conditions. Here, we introduce image-scanning LSM, a method that substantially increases the volumetric imaging speed achievable with LSM while preserving key performance metrics, such as spatial resolution and photon efficiency, as well as accessibility. Our implementation, termed image-scanning oblique plane (ISOP) microscopy, enables volumetric fluorescence imaging at up to 1,000 volumes per second with submicrometer lateral spatial resolution. We demonstrate the broad utility of ISOP microscopy by recording and analyzing the dynamics of behaving and rapidly moving organisms.

## Introduction

Comprehensive three-dimensional observation of structural and physiological dynamics is an effective approach to understanding complex biological systems. Volumetric fluorescence microscopy has enabled such observations across spatial scales ranging from subcellular organelles to whole organisms, thereby advancing fields such as cell biology, developmental biology, and neuroscience (1–6). In particular, light-sheet microscopy (LSM) is a leading platform for practical volumetric fluorescence imaging as it combines low phototoxicity, minimal photobleaching, high spatiotemporal resolution, and low computational burden (7–9). Yet even state-of-the-art LSM remains limited in volumetric imaging speed because optically sectioned images must still be acquired sequentially.

Efforts to improve the imaging speed of volumetric fluorescence microscopy have spanned optical, hardware, and computational strategies, but each entails significant trade-offs that limit practical utility and accessibility (10–18). For example, simultaneous multi-plane imaging approaches employed in light-sheet and widefield microscopy require beam-splitting optics, such as beam splitters and diffraction gratings, which cause significant effective photon loss, thereby compromising the signal-to-noise ratio (SNR) and recording duration (10, 11). A high-speed camera combined with an image intensifier can increase imaging speed while preserving SNR, but its high cost limits accessibility and the intensifier may degrade spatial resolution, depending on the optical design (12). Light-field microscopy enables single-shot volumetric imaging through computational reconstruction, but its spatial resolution is comparable to that of widefield microscopy, which lacks optical sectioning capability. While recent advances have partially mitigated this limitation, it is still better suited for imaging specimens with lower structural complexity or thinner geometries (13–16). As a result, *in vivo* observation of dynamics in organisms under naturalistic, unrestrained conditions, such as freely behaving *Caenorhabditis elegans* (19–21), remains technically limited.

Here, we introduce image-scanning LSM, a method that substantially increases the volumetric imaging speed achievable with LSM while maintaining high spatial resolution, photon efficiency, and accessibility. This is achieved by synchronously scanning the excitation and fluorescence light paths to enable simultaneous multi-plane image acquisition. Specifically, we implemented the method in oblique plane microscopy, a highly versatile form of LSM (12, 22–24). This method, termed image-scanning oblique plane (ISOP) microscopy, increases volumetric imaging speeds by up to an order of magnitude compared with conventional oblique plane microscopy, thereby enabling volumetric fluorescence imaging at up to 1,000 volumes per second (vps) with submicrometer lateral spatial resolution. We demonstrate the broad utility of ISOP microscopy by recording and analyzing the dynamics of behaving and rapidly moving organisms, such as neuronal activity in the head of *C. elegans*, muscle-fiber contractions and calcium signals in a tardigrade, and swimming and flow dynamics in *Chlamydomonas reinhardtii*.

## Results

### Principles of ISOP microscopy

The key idea behind our method for increasing the volumetric imaging speed of LSM is to minimize the number of pixels that are read out from the camera but not used for volumetric image reconstruction. In conventional LSM, a CMOS camera—commonly used in fluorescence microscopy—outputs image data via line-by-line readout. As a result, if the imaging goal requires a field of view of an optically sectioned image that is small relative to the camera’s region of interest (ROI) in the readout direction, the raw data contains unused pixels, thereby lowering the effective image acquisition rate (Fig. 1*A* top). Our method addresses this limitation by filling the ROI with a series of sectioned images at different depths (i.e., focal plane positions), thus fully using the acquired pixels for volumetric image reconstruction and maximizing the effective data readout rate (Fig. 1*A* middle). This multi-plane imaging approach is implemented by incorporating a commonly used optical scanner into the LSM fluorescence imaging system, and is applicable in principle to both selective plane illumination microscopy (SPIM) (7) and digital scanned light-sheet microscopy (DSLM) (9), the two fundamental forms of LSM (Fig. 1*B* and Movie S1). In SPIM, where a sheet-shaped excitation beam illuminates the entire field of view within a focal plane, an image scanner moves the position of the fluorescence image on the camera’s image sensor while a focal plane scanner moves the focal plane and the excitation beam. In addition, a light source stroboscopically modulates the excitation beam to avoid motion blur. Thus, multiple fluorescence images are formed on the image sensor during its exposure time. However, the short illumination time severely limits imaging sensitivity, which is particularly critical in high-speed imaging. By contrast, in DSLM, an axially symmetric excitation beam (e.g., Gaussian beam) is repeatedly scanned across the field of view during image scanning by an excitation beam scanner. In this case, the tightly confined beam effectively excites the sample while avoiding motion blur, allowing for image acquisition with sufficiently high imaging sensitivity (typically more than tenfold higher than in SPIM) using standard laser sources with output powers below 100 mW (*SI Appendix, Supplementary Note S1*). This image acquisition process results in linear distortion of each sectioned image, such as skew (shear distortion), shrinkage, or expansion, depending on the beam scanning direction (Fig. 1*A* bottom), but it is corrected by resampling during volumetric image reconstruction, where the sectioned images are cropped from a series of image frames and stacked (Fig. 1*C*). In contrast to previous multi-plane imaging approaches (10, 11), our method results in minimal fluorescence photon loss due to its simple implementation using a beam scanner. Based on the principles of image scanning microscopy (25), imaging via excitation beam scanning and image scanning leads to changes—mostly decreases—in spatial resolution; however, these changes become negligible when the beam scanning speed is sufficiently higher than the image scanning speed (*SI Appendix, Supplementary Note S2*).

**Fig. 1.**
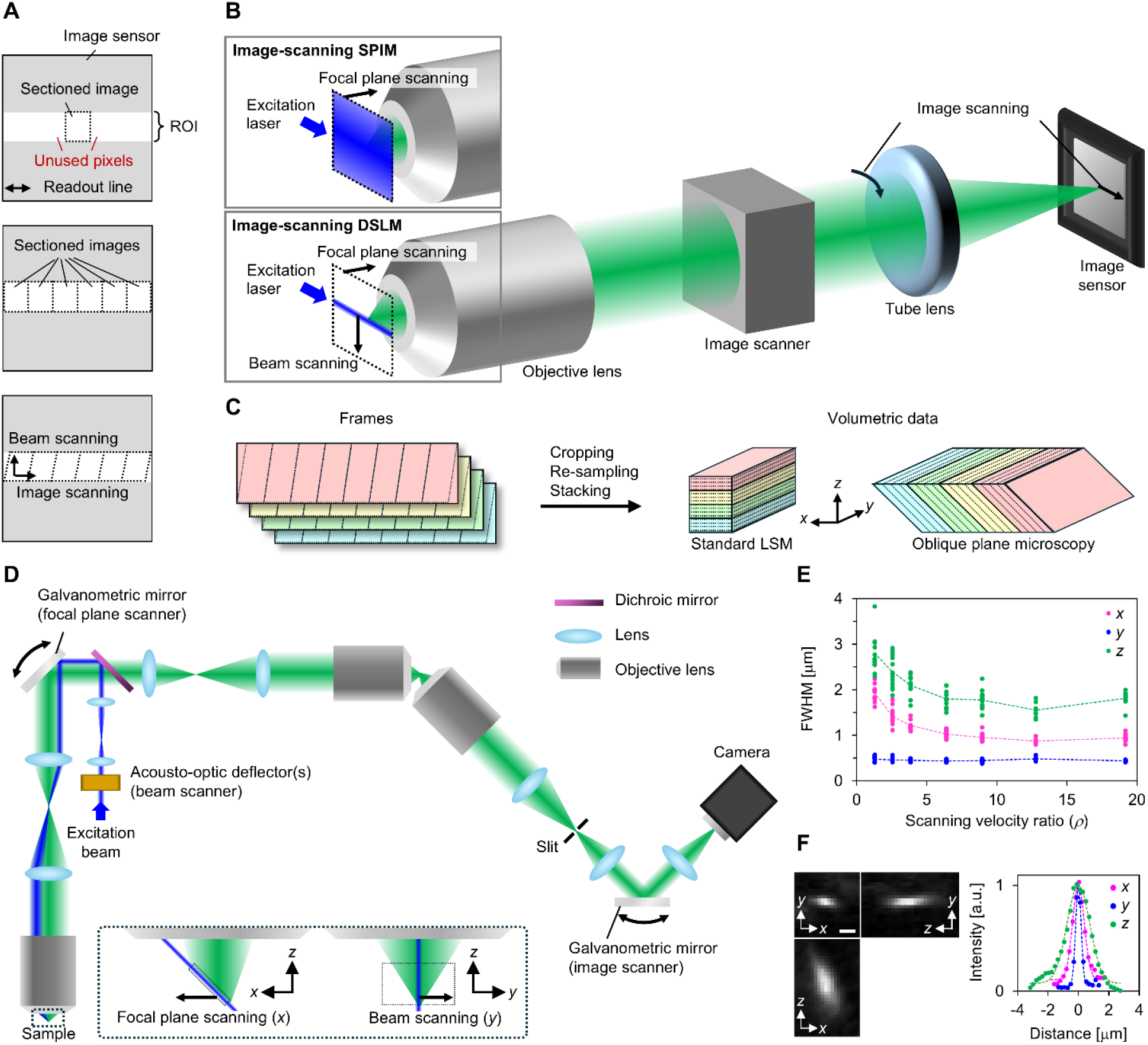
Principles and spatial resolution of ISOP microscopy. (*A*) Image formation on an image sensor in conventional LSM (top), LSM with multi-plane imaging (middle), and image-scanning DSLM (bottom). (*B*) Schematic of image-scanning LSM. (*C*) Volumetric image reconstruction from raw image frames. (*D*) Schematic of ISOP microscopy. The inset represents the geometry in the object space. (*E*) FWHM values of point spread functions (PSFs) along the *x*, *y*, and *z* directions at different scanning velocity ratios (*ρ*), obtained from 95 fluorescent beads. The dashed lines indicate the mean FWHM values at each scanning velocity ratio *ρ*. The definitions of the *x*, *y*, and *z* directions are provided in (*C*) and (*D*). (*F*) Representative PSF cross sections along the *xy*, *xz* and *zy* planes (left) and intensity profiles along the *x*, *y*, and *z* directions (right) obtained by imaging a 200-nm green fluorescent bead at *ρ* = 8.9. The dashed lines represent fitted intensity profiles with an offset Gaussian function. The FWHM values along the *x*, *y*, and *z* directions are, respectively, 957 ± 70, 446 ± 29 and 1772 ± 187 nm (mean ± s.d., *n* = 19 fluorescent beads). Scale bar, 1 μm.

We designed and constructed an ISOP microscope, in which the image scanning approach was implemented in a DSLM-based oblique plane microscope setup (Fig. 1*D* and *SI Appendix,* Fig. S1) (12, 22–24). In this setup, we synchronously controlled two galvanometric scanners for focal plane scanning and image scanning, acousto-optic deflectors for excitation beam scanning, and scientific CMOS (sCMOS) cameras, to capture multi-plane images. The volumetric imaging speed and volumetric field of view were determined by configuration parameters, such as the scanning speeds and ranges and the camera exposure time (see Materials and Methods and *SI Appendix,* Fig. S2). The beam scanning direction was perpendicular to the image scanning direction. We placed a slit aperture at an intermediate image plane as an auxiliary element to restrict the width of each sectioned image. We measured the point spread function (PSF) of the optical system and confirmed that it approached a limiting size as the scanning velocity ratio *ρ*—defined as the beam scanning velocity divided by the image scanning velocity on the image sensors—increased (Fig. 1*E*). This limiting size corresponds to the PSF without image scanning. We also confirmed that the PSF size nearly reached the limit at *ρ* = 8.9, where the full width at half maximum (FWHM) values (957 nm in *x*, 446 nm in *y*, and 1,776 nm in *z*, where *x*, *y*, and *z* are the focal plane scanning, beam scanning, and depth directions, respectively) were consistent with those in a previous report on a similar optical setup using three water-immersion objective lenses (Fig. 1*F*) (26). Since a higher *ρ* shortens the irradiation duration of the excitation light, thereby reducing the imaging sensitivity, we adopted *ρ* = 8.9 for all subsequent biological imaging experiments. Further details on the spatial resolution evaluation are provided in *SI Appendix, Supplementary Note S2*.

### Whole-brain calcium imaging of freely behaving *C. elegans* at 50 vps

Our method combines the optical sectioning capability of LSM with high-speed imaging capability, making it ideally suited for volumetric fluorescence imaging of dense structures exhibiting rapid dynamics. To demonstrate this capability, we recorded and analyzed neuronal activity in the head of freely behaving *C. elegans* using ISOP microscopy. We first prepared transgenic worms expressing tdTomato and GCaMP6f in the nuclei of all neurons and acquired two-color volumetric images of each worm’s heads at 50 vps for 196.2 s (Fig. 2*A*; *SI Appendix,* Fig. S3*A*; and Movies S2 and S3), where each single-color volumetric image was reconstructed from eight consecutive image frames captured at 400 Hz using an sCMOS camera. This volume rate is 4.5 times higher than that expected from the same setup without image scanning (*SI Appendix,* Fig. S2) and approximately 5–35 times higher than those previously reported for the whole-brain imaging of *C. elegans* using conventional methods (e.g., spinning-disk confocal microscopy and oblique plane microscopy) (2, 12, 13, 19–21, 27). An automated stage tracking system ensured that each worm’s head stayed within the lateral field of view of the volumetric imaging during image acquisition (see Materials and Methods and *SI Appendix,* Fig. S4) and recorded its positions over time, from which the trajectory and head movement speed were derived (*SI Appendix,* Fig. S5).

**Fig. 2.**
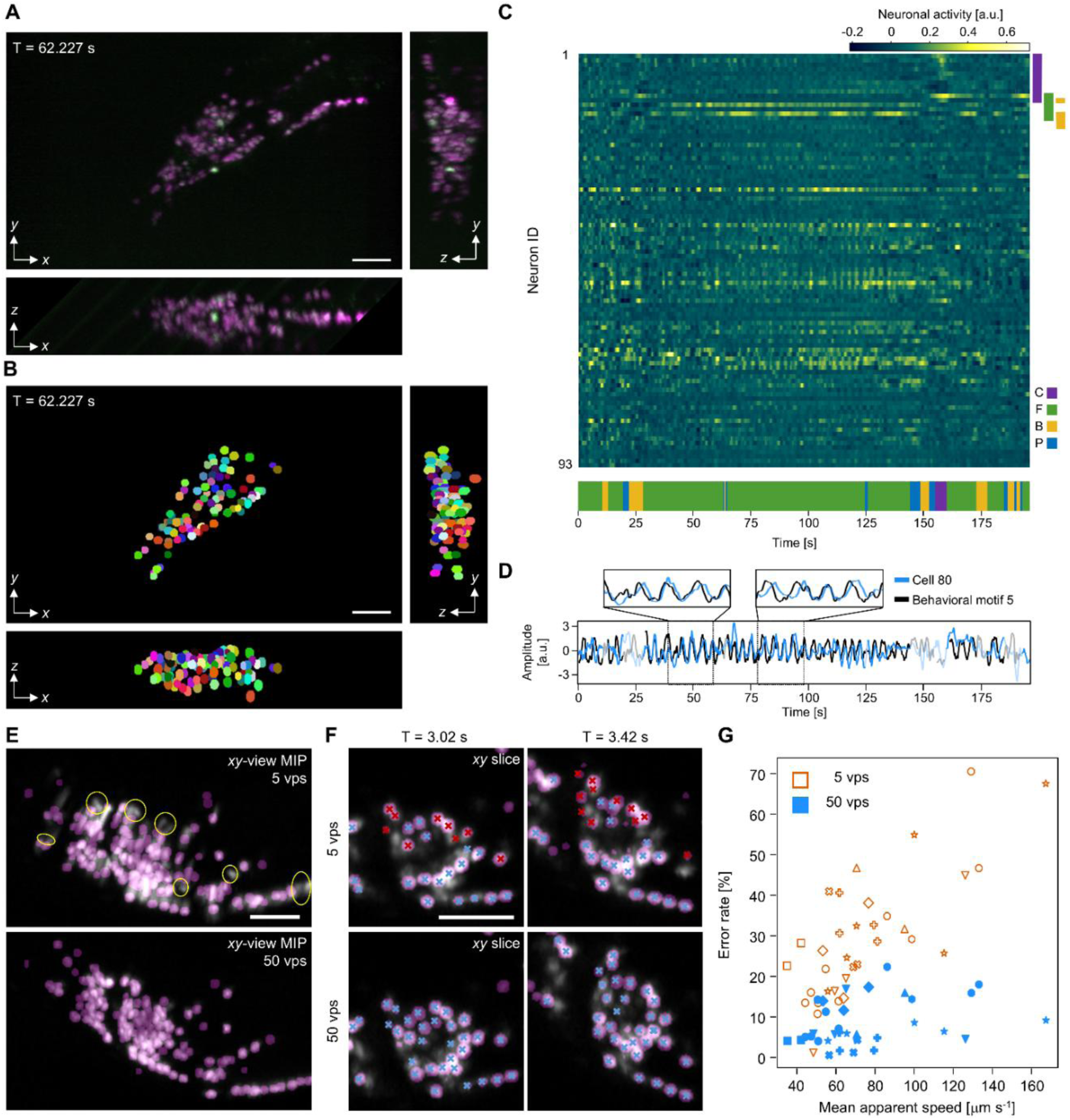
Whole-brain calcium imaging of freely behaving *C. elegans* at 50 vps. (*A*) Maximum intensity projections (MIPs) in the *xy*, *xz*, and *zy* planes of a single volumetric image acquired at 50 vps from the head region of a freely behaving *C. elegans* (Worm 1). Dual-color images show neuronal nuclei labeled with tdTomato (magenta) and GCaMP6f (green). Scale bar, 20 μm. (*B*) Cell tracking results. 93 reliably tracked neurons are labeled with distinct colors. Scale bar, 20 μm. (*C*) Time series of neuronal activity (top) and behavioral states (bottom). The heatmap displays 196.2 seconds of neuronal activity from the 93 reliably tracked cells. Each row corresponds to a neuron and is color-coded to indicate neuronal activity amplitudes. Missing data due to occasional cell tracking failures are shown in magenta. In the time series of behavioral states, manually annotated states are shown with colors denoting coiling (C, purple), forward locomotion (F, green), backward locomotion (B, yellow), and pausing (P, blue). Colored bars on the top right indicate neurons correlated with specific behaviors (|*r*| > 0.3, *r*: Pearson correlation coefficient). (*D*) Selected neuronal activity (blue) and the amplitude of a behavioral motif (black) during forward locomotion. The time series during forward locomotion is highlighted. The insets show enlarged views that highlight the periodicity and correlation of the signals. (*E*) Representative detection errors in tdTomato-labeled neurons in the head of a moving worm. Top-down MIPs of a simulated 5-vps volumetric image (top) and the corresponding original 50-vps volumetric image (bottom) are shown in grayscale, with detected cell regions overlaid in magenta. Detection errors are highlighted by yellow circles. Scale bar, 20 μm. (*F*) Representative tracking errors over time for tdTomato-labeled neurons in the moving head of a worm. *xy* slice images of a simulated 5-vps volumetric image (top) and the corresponding original 50-vps volumetric image (bottom) are shown in grayscale, with detected cell regions overlaid in magenta, at T = 3.02 s (left) and T = 3.42 s (right). Cells successfully and unsuccessfully tracked by a pretrained feedforward neural network (see Materials and Methods) are marked as blue and red crosses, respectively. This judgment was based on visual inspection of the tracking results. Scale bar, 20 μm. (*G*) Tracking error rates (vertical axis) plotted against mean apparent speed of tracked cells (horizontal axis), obtained from 10 datasets (10 organisms). The red and blue points represent 5-vps and 50-vps images, respectively. The different point shapes represent different temporal windows.

We detected and tracked cells in volumetric images from the tdTomato channel and extracted the time-series activity of up to 93 head neurons from both the tdTomato and GCaMP6f channels, while also classifying the behavioral state at each time point by visual inspection of brightfield images (Fig. 2 *B* and *C*; *SI Appendix,* Fig. S3 *B* and *C*; and Movie S4 top).

We examined the relationship between the extracted neuronal activity and behavioral patterns. We first assessed the correlation between each behavioral state and the neural activity of each recorded neuron for a representative worm (Worm 1), revealing correlations (|*r*| > 0.3, *r*: Pearson correlation coefficient, *n* = 1999 time points) between multiple neurons and specific behaviors (Fig. 2*C* top and *SI Appendix,* Fig. S6). In addition, we quantified the behavior of the worm as a time series of posture vectors, each consisting of local angles along the centerline of the worm’s head, and decomposed it into the time series of behavioral motif amplitudes using a variant of independent component analysis (see Materials and Methods and *SI Appendix,* Fig. S7 *A* and *B*) (28). The results showed a neuron associated with a particular behavioral motif corresponding to head swing (Fig. 2*D* and *SI Appendix,* Fig. S7 *C* and *D*). These results demonstrate that ISOP microscopy enables the investigation of the relationship between diverse behavioral repertoires involving rapid movements and neuronal activity of *C. elegans*, making it a powerful tool for studying the neural circuit mechanisms underlying behavior (21, 29).

We further investigated how the volumetric imaging speed affects the accuracy of cell detection and tracking. To this end, we first generated volumetric images by simulating low-speed image acquisition at 5 vps, which is comparable to those in previous whole-brain imaging studies of nematodes, from experimentally acquired images originally obtained at 50 vps, and then performed cell detection and tracking using the same procedure applied to the original images. These simulated images included motion-induced image deformation caused by the relatively long image acquisition time of 200 ms, severely impairing cell detection accuracy (Fig. 2*E*). In addition, a substantial increase in tracking errors was observed in the simulated images, most likely due to cell detection errors and large postural changes between consecutive images (Fig. 2*F*) (30). To precisely compare cell tracking performance between the 5-vps and 50-vps datasets, we evaluated tracking error rates and average neuronal velocities for 34 temporal windows of 400 ms from 10 datasets (10 worms). The results showed that the tracking error rates in the 5-vps dataset significantly increased with the average neuronal velocities, whereas those in the 50-vps dataset remained low across all windows (Fig. 2*G*). Furthermore, for comparison, we acquired volumetric images of the head of a freely behaving worm exhibiting comparable head movement speed using conventional spinning-disk confocal microscopy at 3.2 vps (*SI Appendix,* Fig. S8 *A* and *B*) and performed cell detection and tracking on the acquired images. However, cell detection and tracking performance progressively deteriorated during rapid head movements and eventually became infeasible, during which motion blur, deformation, and large inter-volume cell displacements were frequently observed (*SI Appendix,* Fig. S8 *C* and *D*, and Movie S4 bottom). These results demonstrate that high-speed volumetric imaging with ISOP microscopy enables the robust recording of neuronal activity during rapid head movements in *C. elegans*, outperforming conventional volumetric microscopy used for whole-brain imaging of this organism.

### Volumetric imaging of muscle dynamics in a tardigrade during locomotion at 10 vps

Our method is broadly effective for the quantitative morphological analysis of dynamically changing structures. To demonstrate this capability, we employed ISOP microscopy to observe muscle fibers during locomotion in the transparent tardigrade *Hypsibius exemplaris*, a tractable model organism for studying neuromuscular control and the biomechanics of movement (31). Tardigrades are among the smallest animals capable of coordinated, limb-driven walking; their musculoskeletal system is specialized for locomotor control, offering unique insights into the evolution of movement in multicellular organisms (32). Using a recently developed genetic tool, the TardiVec system (33), we expressed mCherry and GCaMP6s in muscle fibers of *H. exemplaris*. We then acquired two-color volumetric images at 10 vps for 64.8 s during locomotion while tracking the animal using the automated stage tracking system (Fig. 3*A*, *SI Appendix,* Fig. S9, and Movie S5). This volume rate is 1.8 times higher than that achievable using the same setup without image scanning (*SI Appendix,* Fig. S2).

**Fig. 3.**
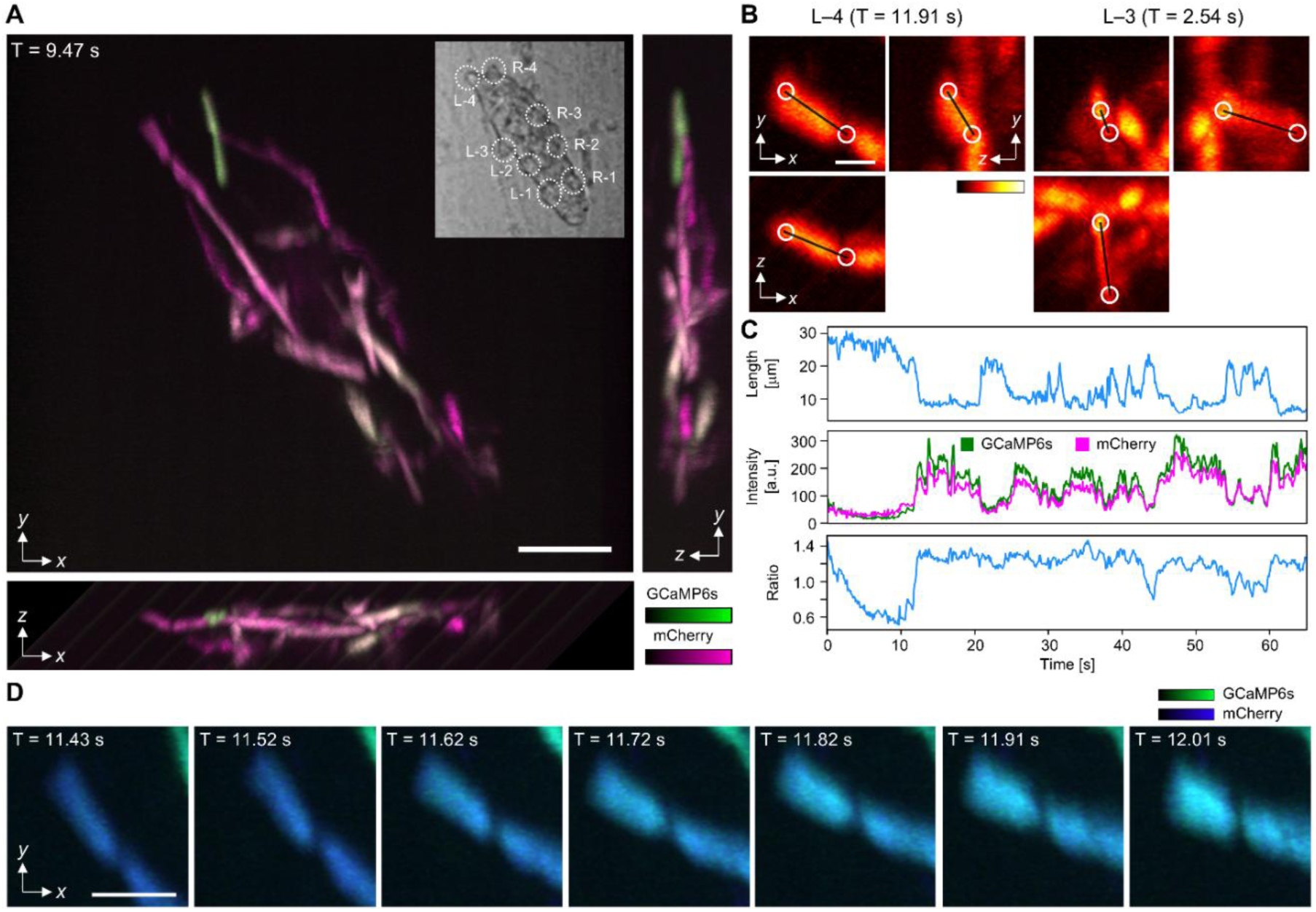
Volumetric imaging of muscle dynamics in a tardigrade during locomotion at 10 vps. (*A*) MIPs in the *xy*, *xz*, and *zy* planes of a volumetric image acquired using ISOP microscopy at 10 vps, showing co-expression of mCherry (red-biased magenta) and GCaMP6s (green) in muscle fibers across the entire body of the *H. exemplaris*. The inset shows a brightfield image of the same animal with adjusted contrast and brightness, with the first to fourth left (L–1 to L–4) and right (R–1 to R–4) legs labeled. Scale bars, 50 μm. (*B*) MIPs of volumetric mCherry fluorescence images of muscle fibers in the fourth left leg (left) and the third left leg (right), illustrating the three-dimensional orientation of muscle fibers. The circles denote the defined regions corresponding to the joint sites with adjacent muscles and the distal ends, respectively. Scale bar, 10 μm. (*C*) Time series measured from a muscle fiber in L–4, showing muscle fiber length (top), GCaMP6s and mCherry fluorescence signals (middle), and ratiometric calcium activity (bottom). (*D*) Time series of top-down MIPs (T = 11.43-12.01 s) of dual-color volumetric images of a muscle fiber in L–4, illustrating both fiber contraction and calcium level changes. Changes in calcium levels are visualized as color changes, reflecting the ratiometric signal. The MIPs were smoothed using a Gaussian filter with a standard deviation of 0.5 pixels. Scale bar, 20 μm.

We analyzed the acquired volumetric images to investigate neuromuscular control during locomotion. Based on anatomical landmarks (34), we identified multiple muscle fibers in the fluorescence images from the mCherry channel. These fibers had various orientations in three dimensions, yet volumetric imaging enabled accurate length measurement (Fig. 3*B*). We measured the length of a representative muscle fiber located in the fourth left leg by tracking two small spherical target regions positioned at its ends (Fig. 3*C* top). Additionally, we calculated the ratio of the total fluorescence intensity of GCaMP6s to that of mCherry within the target regions (Fig. 3*C* middle and bottom). This ratio was used as a proxy for the total amount of calcium ions within the muscle fiber, as it excludes apparent calcium signal changes due to muscle contraction and relaxation. The muscle fiber length and the ratio were negatively correlated (*r* = −0.72, *n* = 665 time points), confirming the presence of calcium-dependent neuromuscular contraction dynamics. This negative correlation was also visually apparent in the images (Fig. 3*D*).

Notably, the high imaging speed of ISOP microscopy not only enables tracking of rapid dynamics across consecutive volumetric images, but also contributes to reducing morphological measurement errors within individual volumetric images. In light-sheet or point-scanning microscopy such as confocal microscopy, where sectioned images are acquired sequentially, object motion during the acquisition of a single volumetric image causes image deformation artifacts, compromising morphological measurement accuracy (*SI Appendix, Supplementary Note S3*). In our experiments, based on the maximum movement speed of the muscle fibers (∼300 μm s^−1^) and the volumetric imaging rate, estimated measurement errors of the muscle fiber length due to deformation artifacts were less than 10%, which would increase by 1.8-fold without image scanning. These errors can be further reduced by estimating the profile of motion-induced image deformation from the estimated instantaneous velocities of the muscle fiber.

### Volumetric imaging of rapidly moving *C. reinhardtii* cells at 1,000 vps

Our method enables ultrafast volumetric imaging at rates exceeding 1 kHz by adjusting the field of view, offering potential to uncover fast biological dynamics that have remained largely unexplored. One such application is the study of microbial motility, which is fundamental to understanding microbial behavior, navigation, and adaptation in diverse ecological environments (35–37). We employed ISOP microscopy to observe the motile unicellular alga *C. reinhardtii*, a versatile model organism that has provided key insights into various biological processes, including photosynthesis (38), flagellar motility (36), and adaptive locomotion (35). We acquired two-color volumetric fluorescence images of swimming *C. reinhardtii* cells with fluorescently labeled nuclei and autofluorescent chloroplasts in a high-salt medium at 1,000 vps for 10 s (Fig. 4*A* and Movie S6). This volume rate is 13 times higher than that achievable using the same setup without image scanning (*SI Appendix,* Fig. S2). We tracked cell motion using the autofluorescent chloroplast images, which exhibited repeated orbital trajectories (Fig. 4*A*). The swimming speed periodically peaked at ∼100 μm s^−1^, during which flagellar beating was inferred to occur (39) (Fig. 4*B*). In addition, the two-color fluorescence imaging allowed us to track cellular orientations based on the relative positions of fluorescence images of nuclei and chloroplasts (Fig. 4*C* and *SI Appendix,* Fig. S10). We further acquired images of *C. reinhardtii* cells rapidly swept by a flow (Movie S7). Despite flow speeds exceeding 1 cm s^−1^, volumetric images of cells with minimal motion-induced deformation artifacts were acquired (*SI Appendix, Supplementary Note S3* and Fig. 4*D*). These results demonstrate that ISOP microscopy is well suited for investigating the motility of a wide range of microorganisms. In contrast to previously reported methods for measuring the 3D motility of single-cell organisms such as stereo imaging (35, 36) and digital holography (40), ISOP microscopy offers more than an order of magnitude higher imaging speed and high-resolution fluorescence-based visualization of intracellular structures. Its high imaging speed comes at the expense of the field of view, limiting the spatial tracking range, but this limitation can be mitigated by automated stage tracking with 3D position control (19, 20).

**Fig. 4.**
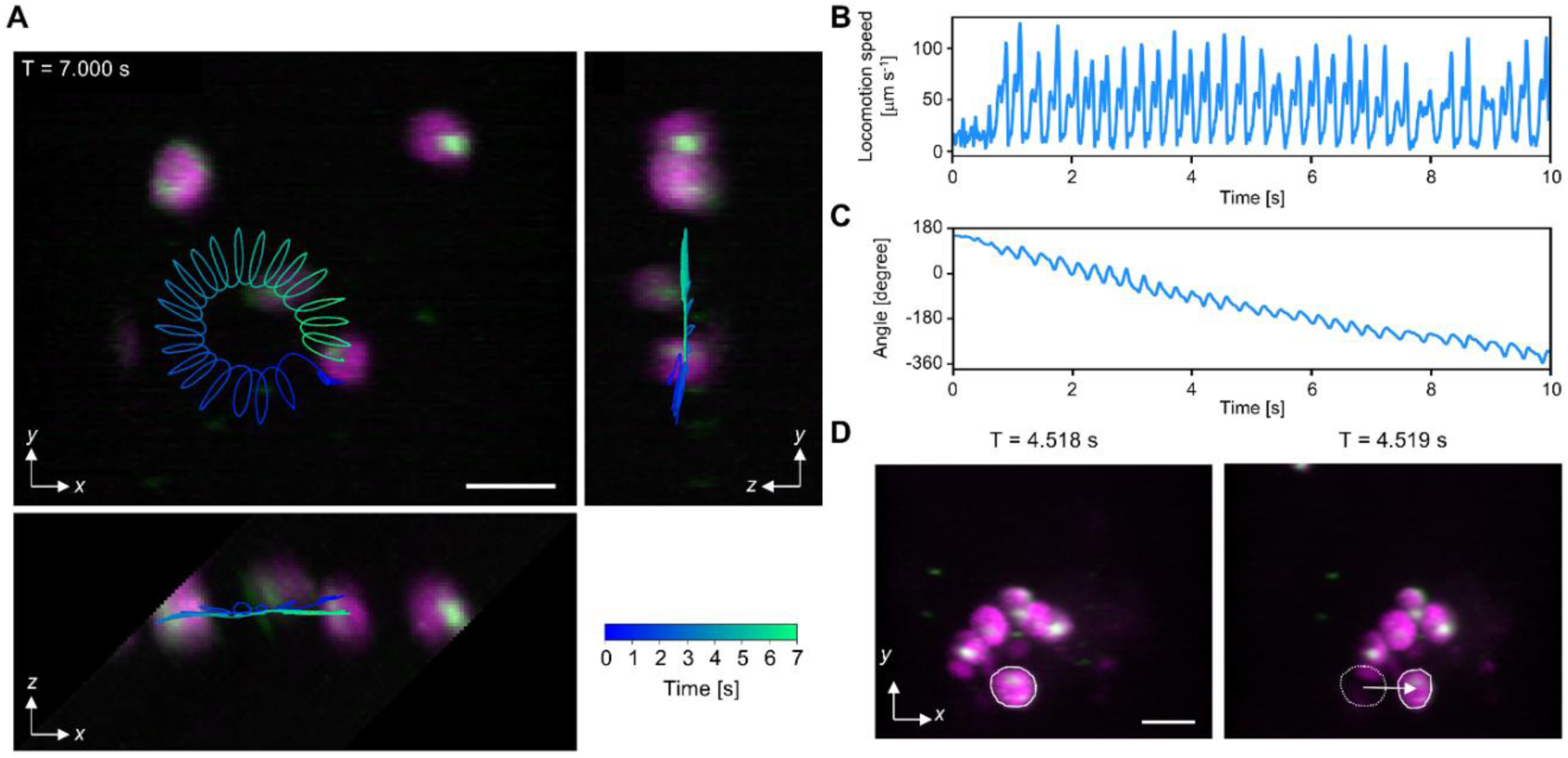
Volumetric imaging of rapidly moving *C. reinhardtii* cells at 1,000 vps. (*A*) MIPs in the *xy*, *xz*, and *zy* planes of a volumetric image of moving *C. reinhardtii* cells acquired using ISOP microscopy at 1,000 vps with autofluorescent chloroplasts (magenta) and SYTO16-stained nuclei (green). The overlaid trajectory shows the 3D displacement of a cell marked by the center of its chloroplast image over a 7-second period, with color indicating temporal progression. Scale bar, 10 μm. (*B*) Time series plot of the locomotion speed of the *C. reinhardtii* cell whose trajectory is shown in (*A*). (*C*) Time series plot of the orientation in the *xy* plane of the same cell. (*D*) Top-down MIPs of flowing *C. reinhardtii* cells captured by ISOP microscopy at T = 4.518 s (left) and T = 4.519 s (right), highlighting a cell that moved 10.65 μm in 1 ms at an instantaneous flow speed of 10.65 mm s^−1^, with autofluorescent chloroplasts (magenta) and SYTO9-stained nuclei (green). Scale bar, 10 μm.

## Discussion

We demonstrated that our image-scanning-based multi-plane imaging method significantly enhances volumetric imaging speeds in LSM while maintaining other performance metrics such as spatial resolution and fluorescence photon efficiency, allowing us to capture high-speed biological dynamics at rates of up to 1,000 vps. Although high-speed fluorescence imaging typically reduces imaging sensitivity or shortens imaging durations due to photobleaching, our results indicate that sufficient imaging speeds, SNRs, and imaging durations can be achieved simultaneously under practical conditions owing to the photon-efficient nature of our method. The method also retains practical accessibility through its simple scanner-based implementation and compatibility with standard sCMOS cameras. As demonstrated by the above experimental results, the method enables flexible software-based adjustment of the field of view without any modification to the optical hardware, while maintaining nearly the maximum effective pixel readout rate dictated by the camera, offering a unique advantage over existing volumetric imaging techniques. We implemented the method in oblique plane microscopy for experimental demonstrations, but it is broadly applicable to other types of LSM, significantly expanding the range of imaging targets for existing light-sheet microscopes.

As demonstrated above, the high-speed volumetric imaging capability of the method enables *in vivo* interrogation of fast neural, muscular, and behavioral dynamics of organisms under naturalistic conditions—biological processes that have remained largely inaccessible with conventional volumetric imaging techniques. Notably, our method achieves both high spatial resolution in three dimensions and exceptionally high imaging speed, making it a uniquely practical option for volumetric imaging of dense, rapidly changing structures, such as the head neurons of a freely behaving nematode. In addition, volumetric imaging at exceptionally high speeds exceeding 100 Hz allows a wide range of applications, such as membrane potential imaging with millisecond temporal changes, imaging flow cytometry, and particle tracking. These capabilities open new opportunities for probing biological events across spatiotemporal scales that have been challenging to access.

Because the method enables faster volumetric imaging by minimizing unused pixels in the readout, it is less effective—or is even ineffective—for large structures whose sectioned images occupy most or all of one dimension of the image sensor or for dim structures that require exposure times substantially exceeding the frame readout time. The effectiveness of the method can be evaluated by theoretical calculations based on the number of voxels in the three dimensions and the required exposure time, allowing for prior verification in an experimental design or preliminary snapshot imaging experiments (see Materials and Methods and *SI Appendix,* Fig. S2). For dim structures for which our method is ineffective, conventional SPIM serves as a primary alternative because it allows fluorescence imaging with continuous excitation, which helps increase fluorescence intensity. Notably, the DSLM-based ISOP microscopy setup can operate in conventional SPIM mode by changing the driving signal for the acousto-optic deflectors used for beam scanning and fixing the position of the image scanner (*SI Appendix, Supplementary Note S4*). Thus, within a single microscope setup, it is possible to instantly switch between the high-speed ISOP microscopy mode and the high-sensitivity conventional SPIM mode by changing the software configuration.

The method is compatible with a wide range of techniques and hardware components that enhance the performance of LSM owing to the simplicity of its implementation, further extending its utility. In particular, the method enables both real-time volumetric image reconstruction through computationally efficient pixel resampling and instant switching of imaging conditions (including imaging modes) via software operation, making it highly compatible with the concept of smart microscopy, where microscopes respond adaptively to the state of observed structures or to user-provided information (41–43). Indeed, using the volumetric image data of *C. elegans* neurons obtained with ISOP microscopy, we confirmed that real-time cell detection and tracking are feasible using a custom-designed computationally efficient algorithm (*SI Appendix,* Fig. S11 and Movie S8). Thus, the method enables experimental designs that were not possible with conventional microscopes, such as adaptive illumination adjustment and real-time targeted optical stimulation (2, 44). In addition, the method is also compatible with spatial resolution enhancement techniques such as DaXi (24) and lattice light-sheet microscopy (43). As demonstrated in this study, since our method enables high-speed volumetric imaging with standard sCMOS cameras, it is economically feasible to increase the number of color channels by adding cameras while maintaining high volumetric imaging speed. When combined with high-speed cameras equipped with image intensifiers, the method can further increase the imaging speed, albeit at a higher cost. Alternatively, the use of newer-generation sCMOS cameras with higher pixel readout rates also leads to improved volumetric imaging speed. Furthermore, our method serves as an alternative to previously reported multi-angle projection imaging (17), which also involves image scanning operations. Multi-angle projection imaging is particularly suitable for quickly previewing the overall volumetric structure of specimens or efficiently acquiring information from sparse specimens through dimension reduction, while our method is suitable for acquiring complete volumetric images from any specimen, especially dense ones. As both methods can be implemented using a common microscope setup, it is possible to switch between them as different imaging modes by modifying software configurations (*SI Appendix, Supplementary Note S4*).

## Materials and Methods

### Optical setup

A detailed optical setup is schematically shown in *SI Appendix,* Fig. S1. Full lists of optical and electronic components and experimental conditions are shown in *SI Appendix,* Tables S1 and S2, respectively. The optical design based on ray tracing simulation is described in *SI Appendix, Supplementary Note S5*. An excitation beam from a laser source (λ = 488 nm, Genesis CX488-2000 STM-SV, Coherent) was scanned by a pair of acousto-optic deflectors (AOD1 and AOD2, TED-150-100-488, Brimrose) and the first galvanometric scanner (G1, GVS211/M, Thorlabs), which then illuminated the sample with an incidence angle of 45° through the first objective lens (OL1, NA = 1.0, water immersion, XLUMPLFLN 20XW, Olympus). The beam was shaped by anamorphic prisms (AP1 to AP4) and a series of lenses (L1 to L6) to effectively cover the active apertures of AOD1 and AOD2 and to have a beam waist radius of 3 μm (*e^−^*^2^ intensity, in all directions) at the object space of OL1. The series of lenses was also used to make the apertures of AOD1, AOD2, and G1, and the entrance pupil of OL1 conjugate with each other. OL1 was also used to collect fluorescence from the sample. An additional pair of lenses (L7 and L8) were placed to make the entrance pupil of the second objective lens (OL2, XLUMPLFLN 20XW) conjugate with that of OL1 and the aperture of G1. A dichroic mirror (DM1, Semrock Di01-R488/561) was placed between G1 and L7 to introduce the excitation beam into the fluorescence light path. The intermediate image created at the object space of OL2 had a magnification of 1, satisfying the aberration-free remote focusing condition (22). The third objective lens (OL3, XLUMPLFLN 20XW) was oriented by 45° with respect to OL2. A water chamber was placed between the horizontally positioned OL2 and OL3 to maintain immersion. More details about the water chamber are described below. A pair of lenses (L9 and L10) was placed to make the entrance pupil of OL3 and the aperture of the second galvanometric scanner (G2, GVS211/M, Thorlabs; Saturn 9B, ScannerMax) conjugate. A slit aperture (VA100, Thorlabs) was placed at an intermediate image plane between L9 and L10 to adjust the width of each sectioned image.

Images of different fluorescence wavelength bands split by a dichroic mirror (DM2, FF580-FDi01, Semrock) were captured by two sCMOS cameras (CAM1 and CAM2, ORCA Fusion C14440-20UP, Hamamatsu) through tube lenses (L11 and L12, TTL100-A, Thorlabs). Additional filters were placed in front of the tube lenses for each experiment. An LED light (λ = 850 nm) also illuminated the sample from the rear side relative to OL1 so that brightfield images of the sample were captured by a CMOS camera (acA2040-120uc, Basler) through OL1, lenses (L13 and L14), and a dichroic mirror (DM3, FF699-FDi01-t3-25×36, Semrock) that transmitted the LED light and reflected the fluorescence and excitation light. The sample position was adjusted either manually or by a custom-built automated stage tracking system (see below). Among possible configurations for the relative orientation between beam scanning and image scanning, we adopted an orthogonal arrangement (Fig. 1*A* bottom, see *SI Appendix, Supplementary Note S6* for alternatives). The laser output was set to 20–90 mW at the source.

### Automated stage tracking system

A custom-built automated stage tracking system was incorporated into the ISOP microscopy setup to track moving organisms such as *C. elegans* and *H. exemplaris* (*SI Appendix,* Fig. S4). The system physically tracked a target position in the sample using an optical flow-based image analysis combined with motorized stage control. The operation began with brightfield image acquisition at 100 Hz using a CMOS camera (acA2040-120uc, Basler). Images were transferred to a computer equipped with a Xeon W-2265 CPU for real-time processing. An object tracking algorithm was implemented using the Lucas–Kanade method via the OpenCV library (function: calcOpticalFlowPyrLK), which computes optical flow, for identifying and continuously tracking a target point. The target point and the center of the microscope’s field of view were defined through a GUI prior to imaging. At approximately 10-ms intervals, the system calculated the displacement between the target point and the field center. Control signals were then transmitted via a PCIe control board (PXEP-CN, Cosmotec) to a stepper motor driver (CTDR-0514ES-2LS, Cosmotec), which actuated a motorized XY stage (OSMS26-50(XY), OptoSigma) to compensate for the displacement. The vertical position of the sample was adjusted manually using a micrometer stage (TSD-653LWP-M6, OptoSigma) and subsequently maintained constant throughout each imaging session. During imaging, the system continuously recorded brightfield images, timestamps, positions of the target region, and XY stage positions. This comprehensive dataset was used for post hoc analysis of locomotion and behavioral dynamics and to validate the performance of the automated stage tracking system.

### Fast beam scanning using acousto-optic deflectors

We used a pair of acousto-optic deflectors (AODs) for beam scanning to cancel the lens effect that occurs within acousto-optic deflectors (45). When the scanning speed of an AOD is high, the frequency gradient across the beam cross-section becomes non-negligible, causing the beam to converge or diverge along the scanning direction (*SI Appendix,* Fig. S12*A*). This results in axial displacement of the focal spot within the scanning plane at the sample. To suppress this effect, we aligned a pair of AODs using a 1:1 relay optical system and drove them with identical signals, so that the focusing and defocusing effects cancel each other (*SI Appendix,* Fig. S12*B*). This dual-AOD configuration effectively mitigates beam displacement. However, scanning with a single AOD remains possible when the scanning is slow or performed at a constant high speed. Under our experimental conditions, assuming a Gaussian excitation beam, the use of a single-AOD configuration was estimated to cause an axial displacement of the beam waist by up to 1.5 times the Rayleigh length at the sample (for *C. elegans* and *C. reinhardtii* imaging), indicating the effectiveness of the dual-AOD configuration. In contrast, under the imaging condition for *H. exemplaris*, the estimated axial shift was approximately 0.5 times the Rayleigh length, suggesting relatively minor influence.

### Water chamber

We employed a custom-built water chamber between OL2 and OL3 to enable low-cost, high-NA implementation of oblique plane microscopy. In contrast to a previous implementation (26), our water chamber enables both compact implementation and fine alignment of the relative position between OL2 and OL3. The chamber consists of a natural rubber tube made from a disposable glove and metal plates that fit on the tapered region near the tip of the objective lenses (*SI Appendix,* Fig. S13 *A* and *B*). Since the chamber and the objective lenses are held in close contact by spring tension, no leakage occurred even if the relative position between the objective lenses is adjusted (*SI Appendix,* Fig. S13 *C* and *D*). The appearance of the chamber is shown in *SI Appendix,* Fig. S13*E*. The chamber was durable for several months, but further extending its lifetime may be possible by replacing the natural rubber with other elastic materials.

### Control system

The control system of our microscope is schematically shown in *SI Appendix,* Fig. S14. Four function generators (Wavestation 2012, 2052, 2022, Teledyne LeCroy; DG400, RIGOL) generate synchronized trigger signals for AODs and sCMOS cameras, as well as driving signals for galvanometric scanners. An arbitrary waveform generator (AWG, PXDAC4800A-DP, Signatec) generates driving signals for the AODs in response to the trigger signals. The driving signals are amplified by an amplifier (ZHL-1-2W+, Mini-Circuits) and split by a power splitter (ZA2CS-251-20WS+, Mini-Circuits) before driving the AODs. The AWG also generates a monitoring signal that includes burst signals synchronized with the driving signals for the AODs, which is monitored by a USB oscilloscope (PicoScope 4824A, Pico Technology). The other trigger signals directly trigger the sCMOS cameras in the rolling shutter mode to start the exposure. Image data are transferred to workstations and saved as binary files, with color channels saved separately. The driving signals are sent to the driver boards of the galvanometric scanners. Monitoring signals from CAM1 and the galvanometric scanner drivers, which respectively indicate the camera exposure duration and real-time scanning positions, are recorded with the USB oscilloscope. All control processes were implemented in custom-developed LabVIEW programs.

### Image acquisition sequences

In this study, we employed two types of image acquisition sequences (type I and type II), which are schematically illustrated in *SI Appendix,* Fig. S15. In the first one (type I, *SI Appendix,* Fig. S15*A*), the exposure times of the sCMOS cameras are synchronized with a series of beam scans, a linear image scanning in one direction, and a linear focal plane scanning in one direction, while the positions of the image scanner and the focal plane scanner are adjusted during the readout time of the cameras. The positions of the focal plane scanner at the timings of the beam scanning are evenly spaced. An identical waveform is used for the image scanning during each exposure-readout period to ensure that sectioned images are acquired at fixed positions. In addition, to avoid nonlinear distortions arising from the limited response speed of the scanner, the camera’s exposure time is set to fall within a portion of the linear scanning periods of both the focal plane scanning and image scanning. In the second one (type II, *SI Appendix,* Fig. S15*B*), the exposure times of the sCMOS cameras are synchronized with the linear scans in both directions of the focal plane scanner and image scanner. This sequence is better suited for smaller volume imaging, where the required readout time is shorter. We employed type I for imaging of *C. elegans* and *H. exemplaris*, and type II for imaging of *C. reinhardtii*. A theoretical comparison of imaging speeds with these sequences is presented below.

### Imaging speed comparison

The imaging speeds of conventional and image-scanning LSM are governed by the scanner and camera configuration parameters, as well as the imaging field of view, expressed in voxel counts. The volume periods for conventional LSM are given by:

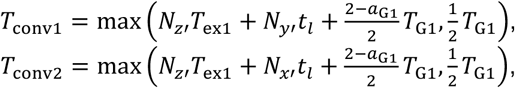

where 𝑁_𝑥′_ and 𝑁_𝑦′_ are the numbers of pixels along the excitation beam propagation direction and its perpendicular direction on the image sensor, respectively, 𝑁_𝑧′_ is the number of sectioned images constituting a single volumetric image, 𝑇_ex1_ is the camera’s exposure time per readout line, which must be greater than the readout time per frame (𝑁_𝑦′_𝑡_𝑙_ for 𝑇_conv1_ and 𝑁_𝑥′_𝑡_𝑙_ for 𝑇_conv2_), 𝑡_𝑙_ is the camera’s line readout time, 𝑇_G1_ is the minimum period of the focal plane scanner, and 𝑎_G1_ is the ratio of linear scanning time to the full unidirectional scanning time of the focal plane scanner. 𝑇_conv1_ and 𝑇_conv2_ denote the volume periods for which the sectioned images are aligned so that the propagation direction of the excitation beam is perpendicular and parallel to the readout direction, respectively. We assumed the rolling shutter mode of the camera. In contrast, the volume acquisition periods of image-scanning LSM are given by:

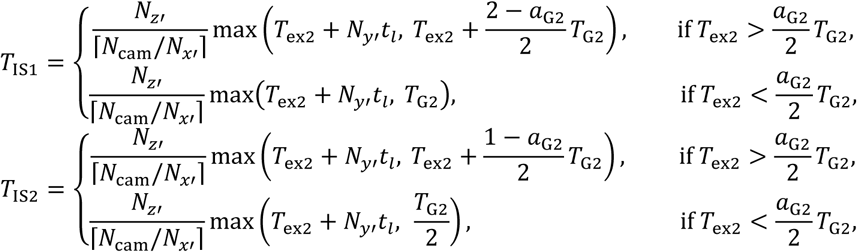

where 𝑁_cam_ is the number of pixels in the camera’s readout line, 𝑇_ex2_ is the camera’s global exposure time, 𝑇_G2_ is the minimum period of the image scanner, and 𝑎_G2_ is the ratio of linear scanning time to the full unidirectional scanning time of the image scanner. 𝑇_IS1_ and 𝑇_IS2_ represent volume acquisition periods for readout sequence types I and II, respectively. For simplicity, the beam scanning speed in DSLM was assumed to be infinitely high. The excitation beam propagation direction was assumed to be parallel with the readout direction on the image sensor. *SI Appendix,* Figure S2*A* shows how these volume acquisition periods vary with 𝑁_𝑦′_. The reduction in volume acquisition periods (i.e., the improvement in imaging speed) becomes particularly prominent for intermediate values of 𝑁_𝑦′_. The volume rates of each imaging mode (𝐹_conv_ ≡ 1/ min(𝑇_conv1_, 𝑇_conv2_) , 𝐹_IS1_ ≡ 1/𝑇_IS1_, 𝐹_IS2_ ≡ 1/𝑇_IS2_) under parameters used for imaging *C. elegans*, *H. exemplaris*, and *C. reinhardtii*, as well as the volume rate enhancement factor achieved by image scanning (𝑔_IS1_ ≡ min(𝑇_conv1_, 𝑇_conv2_) /𝑇_IS1_, 𝑔_IS2_ ≡ min(𝑇_conv1_, 𝑇_conv2_) /𝑇_IS2_) and the parameters used, are shown in *SI Appendix,* Fig. S2*B*. We assumed that 𝑇_ex1_ is equal to the readout time per frame for conventional LSM, ensuring that the minimum possible volume acquisition period is achieved for each 𝑁_𝑦′_. As illustrated in the plots, a wide practical range of 𝑁_𝑥′_ and 𝑁_𝑦′_ values, spanning from approximately 100 to several hundred, shows substantial imaging speed enhancement. Under conditions with relatively small 𝑁_𝑥′_ and 𝑁_𝑦′_ values—such as those used for imaging *C. reinhardtii* cells—a greater than tenfold increase in imaging speed can be achieved.

### Calibration for image acquisition sequences and volumetric image construction

All calibration and volumetric image construction processes were implemented in custom-developed LabVIEW programs.

*Focal plane scanning calibration*.

The calibration of the driving signal for focal plane scanning is schematically illustrated in *SI Appendix,* Fig. S16*A*. First, the monitoring signal from the focal plane scanner—driven by a waveform designed to achieve the target scanning range and sectioned image spacing—and the signal indicating the timings of beam scanning are recorded simultaneously. Then, the voltage values of the monitoring signal corresponding to the timings of beam scanning are recorded.

Here, linear fitting is performed on the dataset of sectioned image indices and their corresponding voltage values in each imaging frame, and the slope and offset of each ramp section of the driving signal are adjusted to compensate for differences from the target linear relation. This is repeated a few times to ensure that the sectioned image spacing reaches the desired constant value.

### Image reconstruction calibration

The workflow for constructing a volumetric image from acquired image frames is outlined in *SI Appendix,* Fig. S16*B*. Initially, each sectioned image is cropped from one of the image frames, and a de-skewing process is applied to correct image distortions caused by beam scanning, followed by intensity correction for each sectioned image. The cropping positions, de-skewing factors, and intensity correction factors are determined via either static or dynamic calibration processes described below. Subsequently, sectioned images optionally undergo vibration removal before being resampled onto the object coordinate system to generate a volumetric image.

### Static calibration

*SI Appendix,* Fig. S16*C* illustrates the static calibration procedure. Without performing focal plane scanning, sectioned images of a fixed calibration sample that includes confluent 10-µm fluorescent beads arranged on a plane are acquired. From these, a representative sectioned image is extracted from the image frames using a specified cropping position in the readout direction and de-skewing factor, and it is designated as the reference image. Thereafter, sectioned images are extracted using preassigned cropping positions, de-skewing factors, and intensity correction factors, and the differences from the reference image are evaluated by the mean square error. These parameters are optimized using the simplex algorithm to minimize the error. If the image scanning direction slightly deviates from the readout direction, vertical offsets in the cropping positions are employed. Additionally, the cropping positions along the readout direction are manually adjusted as needed by reviewing reconstructed images. This process determines the cropping parameters for extracting sectioned images from raw image frames.

### Dynamic calibration

The dynamic calibration includes pre-calibration steps to ensure accuracy under high-speed imaging conditions (*SI Appendix,* Fig. S16*D*). First, calibration images are acquired following the same procedure used in the static calibration, while recording both the monitoring signal of the image scanner and the beam scanning timing signal. Using the acquired dataset, cropping parameters—including cropping positions along the readout line and the de-skewing factor—are determined using the same processing steps as for static calibration. Next, the relationship between cropping positions and monitoring voltages, as well as that between the de-skewing factors and the rates of change of the monitoring voltage, are plotted, and polynomial regression is used to generate calibration curves. During imaging, the monitoring signal from the image scanner and the beam scanning timing signal are continuously recorded. For each sectioned image, the cropping position along the readout line and de-skewing parameter are computed from the calibration curves, recorded monitoring signals, and their derivatives. As in the static calibration, manual adjustment of the cropping position is performed as needed based on inspection of reconstructed images. This dynamic calibration is especially useful at near-maximum scanning speeds of galvanometric scanners, where operational instability may occur. It was specifically applied during *C. elegans* imaging, where such high-speed conditions were used. For all other imaging experiments, static calibration was sufficient and therefore employed.

### Vibration removal

A vibration removal algorithm was employed for the evaluation of point spread functions (*SI Appendix,* Fig. S17*A*). A digital stack of sectioned images cropped from an image frame is first processed to generate a maximum intensity projection (MIP) in the *x’z*’ plane, where *x*′ represents the readout direction, and *z*’ represents the stacking direction. The displacements between adjacent *x*’-direction profiles along the *z*’ axis in the MIP are then computed. Under vibration-free conditions with sufficiently fine spacing along the *z*’ axis, these displacements remain nearly constant. Thus, the average displacement value is considered the true constant, and deviations from this value are identified as vibration components in the *x*’ direction. These deviations are canceled by shifting each *x*′*y*′ slice in the *x*′ direction accordingly, yielding a corrected volumetric image. The same correction process is subsequently applied to a *y’z*’-plane MIP, ensuring the removal of vibration-induced displacements across all *x’y*’ slices. In this process, vibration components perpendicular to the focal plane can cause image artifacts that manifest as non-smooth profiles in the *z*′ direction and may impair accurate PSF size estimation. However, such profiles were rarely observed in practice. We therefore conducted PSF evaluation assuming that the impact of such vibration components was negligible. *SI Appendix,* Fig. S17*B* illustrates a result, showing *x’z*’-plane MIP images of a sample containing 200-nm fluorescent beads before and after the correction.

### Virtual z-position correction

The workflow for virtual *z*-position correction, employed in the analysis of *C. elegans* image data, is illustrated in *SI Appendix,* Fig. S18. This method is applicable when the imaged object has a constrained extent along the *z* direction and exhibits temporal displacement along this axis. To accommodate such displacement, the slit aperture is slightly widened to allow partial overlap of adjacent sectioned images along the readout direction on the camera frame. Under this condition, a reconstructed volumetric image contains shared regions at the upper and lower bounds of the *z* axis in the constructed volumetric image. The volumetric image is projected onto the *z* axis to generate a line profile, which is used to determine the object’s position and spatial extent along the *z* direction. This enables cropping of the *z* range of interest and generation of the final image that does not include the shared region. When the object shifts along the *z* direction, the cropping range is dynamically adjusted to follow the displacement, thereby achieving virtual *z*-position correction. To suppress abrupt displacement caused by estimation errors, the difference in the displacement between consecutive time points is limited to within one voxel. In practice, the *z* displacement of *C. elegans* was gradual, allowing effective tracking of the object’s *z*-axis motion over time.

### Whole-brain calcium imaging of freely behaving *C. elegans* at 50 vps

#### Sample preparation

We used young adult *C. elegans* expressing NLS3-tdTomato and NLS2-GCaMP6f, both driven by the H20p promoter. To minimize behavioral responses to blue light irradiation, we used mutants *carrying* null alleles of *lite-1* and *gur-3*. Worms were cultured on nematode growth medium plates seeded with *E. coli* OP50 at 20 °C in an incubator (CN-25C, Mitsubishi Electric). Sample preparation began by creating an agarose pad (approximately 26 × 26 mm, 3.4% in M9 buffer, NuSieve GTG agarose, Lonza) on a microscope slide, following an established protocol (46).

Worms exhibiting strong tdTomato fluorescence were selected using a fluorescence stereomicroscope (MVX10-LED-4-SW, Evident) and transferred with a platinum picker to an agarose plate without *E. coli*. They were incubated for approximately 30 minutes to reduce residual bacteria attached to the body surface. For imaging, a single worm was transferred to the center of the agarose pad with ∼5 μL of M9 buffer to allow locomotion. A PTFE-PFA film (25-μm thick, NR5100-001, Flon Chemical) was gently placed over the sample to minimize mechanical constraint while permitting free locomotion. The volume of M9 buffer was adjusted as necessary to prevent the worm from floating. The mobility of the worm was confirmed using a stereomicroscope. The microscope slide was then mounted on the ISOP microscope sample holder and fixed at the edges with modeling clay. Distilled water (at room temperature) was added between the objective lens and the film. The worm’s position was manually adjusted, after which the automated stage tracking system was activated.

#### Cell detection and tracking in C. elegans whole-brain imaging data

The following procedure was performed on a modified version of 3DeeCellTracker, a deep learning-based pipeline for segmenting and tracking cells in time lapse volumetric images (47).

Cell detection was performed using individually trained 3D StarDist models for each worm. At four selected volumetric images (1^st^, 3000^th^, 6000^th^, and 9995^th^), we created training datasets, each consisting of the original volumetric image and manually segmented neuronal regions.

Annotations were conducted using ITK-SNAP (v4.2.0). A StarDist model was trained independently on each dataset, with each training session requiring approximately 2 hours. The optimal model for each worm was selected based on two criteria: maximizing the number of neurons detected and minimizing the discrepancy in neuron counts relative to manual annotations. For Worm 1, the model trained on the 6000^th^ volumetric image best satisfied these criteria and was used in all subsequent analyses (*SI Appendix,* Table S3).

Cell tracking was performed using the cell detection results from each volumetric image. We estimated cell correspondences between volumetric images using a pretrained feedforward neural network and then predicted the positions of individual cells in subsequent volumetric images. Specifically, for the original 50-vps datasets, we employed the ensemble tracking mode (47), in which multiple predictions based on correspondences from several preceding volumetric images were averaged to enhance robustness against errors in cell annotations and correspondence estimation. By contrast, for the simulated low-speed imaging datasets at 5 vps, we employed the single tracking mode, in which correspondences between adjacent volumetric images were used for one-step predictions. This choice was made because large inter-volume posture changes in the low-speed images reduce the effectiveness of the averaging operation.

Manual correction was applied to occasional tracking failures due to cells leaving the field of view, significant image deformation, or inconsistent trajectories. To complete the correction within a practical timeframe, we applied it to a subsampled set of volumetric images (10 vps for Worm 1 and 5 vps for Worms 2–4). Corrections were performed using a custom GUI-based tool that enabled efficient visual inspection of tracking results overlaid on raw images.

#### Extracting neuronal signals from volumetric images

Fluorescence intensities from two image channels were extracted for each annotated cell based on the volumetric cell regions identified during cell detection and tracking. For each cell, the mean intensity of voxels within its volumetric region was used as the representative fluorescence intensity for each channel. For both channels, small gaps in the time series of fluorescence intensity due to occasional tracking failures were linearly interpolated. Each interpolated time series was then smoothed using a Savitzky–Golay filter (window size, 1.2 s; polynomial order, 1). To correct for intensity decay due to photobleaching, each time series 𝐼(𝑡) was fitted with a multi-exponential function of the form 𝐼_f_(𝑡) = 𝑎_1_𝑒^−𝑏1𝑡^ + 𝑎_2_𝑒^−𝑏2𝑡^, where 𝑎_1_, 𝑎_2_, 𝑏_1_, and 𝑏_2_ are constants obtained by curve fitting. The photobleaching-corrected signal 𝐼_c_(𝑡) was then defined as 𝐼_c_(𝑡) = 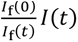. This correction was applied separately to the tdTomato and GCaMP6f channels. The 𝐼_f_(𝑡) ratio of the corrected GCaMP6f intensity to the corrected tdTomato intensity, denoted as 𝑅, was then computed. The neuronal signal was expressed as the fractional change in this ratio, (𝑅 − 𝑅_0_)/𝑅_0_, written as Δ𝑅/𝑅_0_, where 𝑅_0_ was defined as the 10^th^ percentile of the fluorescence ratio time series for each neuron. Finally, the interpolated values corresponding to the original gaps were reverted to missing values (NaNs).

#### Quantitative analysis of worm’s behavior

We quantified the worm’s behavior by representing its head posture at each time point as a vector. To this end, we applied a variance filter twice to each brightfield image and binarized the result using the mean intensity as the threshold. A midline was then extracted via a thinning algorithm and subsequently smoothed using a spline curve. The midline was approximated by 99 consecutive lines of equal length, and the angles of these lines relative to the horizontal axis were computed. From each of the 99 angles, their average was subtracted, yielding a 99-dimensional angle vector representing the worm’s posture in each frame. The time series of these vectors represents the posture dynamics of the worm’s head over time. We computed these vectors for brightfield image frames corresponding to the time points analyzed for tracking, sampled at 10-ms intervals.

We applied time-delay embedded reconstruction independent component analysis (TDE-RICA) (28) to the time series of the angle vector to extract interpretable low-dimensional components. Following a previous study on whole-body posture analysis (48), we used a delay of 8 frames (0.8 s) and 7 independent components. Let 𝑨(𝑡) ∈ ℝ^1×99^ denote the angle vector at time 𝑡. The time-delay embedded vector at time 𝑡 was defined as 𝒙(𝑡) = [𝑨(𝑡 − 7), 𝑨(𝑡 − 6), . . . , 𝑨(𝑡)]ᵀ ∈ ℝ^(99×8)×1^. The resulting time series matrix 𝑿(𝑡) = [𝒙(8), 𝒙(9), . . . , 𝒙(2000)] ∈ ℝ^(99×8)×1992^ was decomposed into a spatial mode matrix 𝑴 ∈ ℝ^7×(99×8)^ and a temporal activation matrix 𝑾 ∈ ℝ^1992×7^ using RICA. The algorithm minimizes the cost function ‖𝑾𝑴 − 𝑿ᵀ‖₂² + ‖𝑔(𝑴)‖₁, where 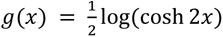 is an entrywise contrast function promoting sparsity. Each row of 𝑴 corresponds to an independent spatial component and can be interpreted as a characteristic behavioral motif. Among the extracted components, the 5th and 6th consistently captured head bending movements.

#### Generation of simulated low-speed imaging data

The method for generating simulated low-speed imaging datasets of *C. elegans* is illustrated in *SI Appendix,* Fig. S19. MIPs in the *xy* plane are generated from the first ten volumetric images acquired at 50 vps. In the projection image at the initial time point, a rectangular region that encloses the worm’s head is manually specified. Corresponding parallelogram-shaped regions in subsequent projection images are then identified as regions that best match the previously defined region under affine transformations. These regions are subsequently used to compute affine transformation parameters that describe changes in the nematode’s posture over time. To interpolate the posture at arbitrary time points, cubic Hermite interpolation is applied to the time series of the four corner coordinates of the regions, allowing us to calculate affine transformation parameters at any desired time point. To simulate depth scanning as performed by a spinning-disk confocal microscope, we assumed that *xy*-plane images at different depths are sequentially acquired at distinct time points. The affine transformations corresponding to the intended time points are applied to the *xy*-plane images at each depth within the reconstructed volumetric dataset. The transformed *xy*-plane images are then stacked to generate a volumetric image that simulates low-speed *z* scanning and includes shear distortions arising from the worm’s motion. Applying this deformation process at 200-ms intervals to each volumetric image produced a time series of virtual low-speed volumetric images, mimicking image acquisition at 5 vps with slow *z* scanning.

#### Evaluation of cell tracking error rate and mean apparent speed of cell movement

The cell tracking error rate and mean apparent cell movement speed were defined for each local temporal window of volumetric images. To compute these metrics, cell tracking was first performed using 3DeeCellTracker on both the original high-speed and simulated low-speed volumetric image datasets. For each volumetric image dataset, a 3D StarDist model was trained independently. A total of 34 local temporal windows, each spanning 400 ms, were extracted from volumetric images of ten freely moving *C. elegans*. Each window contained 21 volumetric images from the original dataset and two volumetric images from the corresponding low-speed dataset. Following cell tracking, the number of successfully tracked cells in each region was manually counted using Napari, a Python-based multidimensional image analysis tool. The tracking error rate was computed as the ratio of cells that failed to be tracked to the total number of detected cells within each temporal window. For each temporal window, the mean apparent speed of cell movement was defined as the average displacement per unit time of all successfully tracked cells across the 21 volumetric images from the original dataset.

#### Analysis of volumetric images acquired by spinning-disk confocal microscopy

The spinning-disk confocal microscope system used in this study has been described previously (49). Briefly, it consisted of an inverted microscope (IX-83, Olympus) equipped with a spinning-disk confocal scanner (CSU-W1, Yokogawa), two sCMOS cameras (ORCA-Flash4.0 v3, Hamamatsu Photonics), a motorized stage (HV-STU1000-T001-A, HawkVision), and custom-built upright imaging optics for monitoring animals on the stage. During free locomotion, tdTomato fluorescence was monitored through the upright imaging optics using a low-magnification objective lens (PLN10X, Olympus) and a CMOS camera (FL3-U3-13Y3M-C, FLIR). Real-time image processing and tracking control were implemented via custom-developed software, which provided closed-loop feedback to the motorized stage, ensuring continuous positioning of the worm’s head within the imaging field of view, in the same manner as the automated stage tracking system implemented in the ISOP microscopy setup. For imaging, individual worms were mounted on a thin agarose pad placed on a 70 × 50 mm coverslip (C050701, Matsunami) and were gently overlaid by a transparent PTFE-PFA film (NR5100, Flon Chemical) to minimize mechanical constraint while permitting free locomotion. Fluorescence images from GCaMP6f and tdTomato expressed in head neurons were acquired using a water-immersion objective lens (LUMPLFLN60XW, Evident) combined with a piezoelectric focus scanner (P-725K085, Physik Instrumente) for volumetric scanning. Images were acquired at a pixel size of 0.43 μm, with each volumetric image composed of 21 optically sectioned images spaced at 2-μm intervals, covering a 40-μm axial range. The first two slices in each volumetric image were excluded from downstream analysis to avoid the impact of image artifacts resulting from the instability of the piezo focus scanner immediately after axial repositioning. Thus, each volumetric image effectively contained 19 planes, each of which consists of 512 × 512 pixels. Volumetric imaging was conducted at 3.2 vps (15 ms per plane), and 1000 volumetric images were collected per imaging session. Image analysis was performed using 3DeeCellTracker, as in the analysis of ISOP microscopy data. Manual segmentation was applied to the first volumetric image in each dataset, which was used for training a StarDist-based model.

#### Real-time cell tracking

We used a custom Python-based cell tracking program to achieve millisecond-scale processing times on a GPU-equipped workstation (NVIDIA RTX A5000 GPU, Intel Xeon W5-3453X CPU). The algorithm consisted of three sequential steps: real-time 3D image reconstruction, initial cell detection, and iterative tracking of identified cells, which required approximately 12 ms, 24 ms, and 4 ms per step, respectively. Volumetric images of tdTomato fluorescence were reconstructed from image frames using the dynamic calibration. Cell positions in the first volumetric image were identified by applying a local maximum filter with a kernel size of 15 × 15 × 15 voxels.

Fluorescence intensity maxima below a predefined threshold were excluded as false positives. Multiple candidates sufficiently distant from the image boundary were selected as tracking targets. In each subsequent volumetric image, cell positions were updated by searching neighborhoods to identify the best-matching voxels. Specifically, for each cell, voxels within a 31 × 21 × 9 voxel neighborhood centered at the previous position were first selected if their absolute intensity difference from the voxel intensity at the previous cell position exceeded a predefined threshold. The cell position was then updated to the intensity centroid computed by applying the watershed algorithm within a 31 × 21 × 9 voxel neighborhood centered at the selected voxel. All processes were performed offline using previously recorded image frames and precomputed cropping parameters.

### Volumetric imaging of muscle dynamics in a tardigrade during locomotion at 10 vps

#### Sample preparation

We used the Z151 strain of *H. exemplaris*, maintained on water-layered agar plates and fed *Chlorella vulgaris* (Chlorella Industry). Muscle fiber-specific expression of mCherry and GCaMP6s was achieved using the TardiVec system, following previously reported microinjection and electroporation protocols with minor modifications for this study (33). The TardiVec system comprises the *H. exemplaris* actin promoter, mCherry or GCaMP6s coding sequences, the 3′ UTR, a selection marker, and a replication origin for *E. coli* amplification. The promoter and 3′ UTR correspond to ∼1 kbp regions flanking an open reading frame (33). For this study, we used a muscle-specific promoter sequence. Plasmid preparation followed the original protocol. For plasmid delivery, glass capillaries (GD-1, NARISHIGE) were pulled using a puller (PC-100, NARISHIGE) at 66.2 °C and 62.0 °C. Microinjections were conducted under an inverted microscope (AXIO Vert.A1, Zeiss) equipped with an injector (IM-31, NARISHIGE) and micromanipulators (MMN-1 and MHW-103, NARISHIGE). After microinjection, individual tardigrades were transferred to distilled water containing algal food. A fluorescence stereomicroscope (MVX10-LED-4-SW, Evident) was used to identify individuals expressing fluorescent proteins in multiple muscle fibers. These animals (six in total) were collected using a 2–200-μL pipette tip and transferred to 2% agarose plates with distilled water but no algal food, and stored in a refrigerator overnight. By the following day, the algal content in their digestive organs had diminished, improving optical clarity during imaging.

To ensure stable positioning during imaging, a microscope slide was lightly scratched with sandpaper to increase surface friction. Two pieces of PTFE-PFA film (50-μm thick, NR5100-002, Flon Chemical) were placed at opposite ends of the slide as spacers. A coverslip was positioned over the film edges and gently pressed down to eliminate air bubbles and to create an enclosed chamber. This gentle compression provided mild restraint that limited the locomotion speed.

Under these conditions, the tardigrades typically adopted one of two orientations: with either the ventral or dorsal side facing the coverslip. Volumetric imaging was feasible in both configurations. The animal shown in Fig. 3 was imaged in the former orientation. Room-temperature distilled water was applied on the coverslip as the immersion medium for the objective lens. Brightfield imaging was used to locate the animals, followed by activation of the automated stage tracking system to maintain the animal at the center of the field of view and in focus. The stage tracking system recorded the animal’s position, which was used to calculate the locomotion speed (see *SI Appendix,* Fig. S9).

#### Analysis of muscle fiber length and calcium signal in a tardigrade leg

To analyze muscle dynamics in a tardigrade, we focused on muscle fibers in the legs identified based on anatomical landmarks (34). For each fiber, two spherical target regions with a diameter of 8.19 μm were defined in the mCherry channel—one at the joint with the adjacent muscle and the other at the distal end—and the same regions were used in the GCaMP6s channel to extract fluorescence intensity values. These regions were manually tracked using Napari. Local calcium levels were computed as the ratio of the average GCaMP6s intensity to the average mCherry intensity within the distal region. Muscle fiber length and fiber tip speed were defined as the distance between the two regions and the displacement of the distal end position per unit time, respectively.

### Volumetric imaging of rapidly moving C. reinhardtii cells at 1,000 vps

*Sample preparation*.

*C*. *reinhardtii* strain 137c cells were cultured in a high-salt medium (50), with a working volume of 20 mL in 50-mL polystyrene suspension culture flasks with filter caps (Greiner Bio-One). Cultures were maintained under static conditions at 28°C with a 14-h light/10-h dark cycle (∼100 mmol m⁻² s⁻¹). Cells were cultured for at least three days before imaging to ensure sufficient biomass. Cells were harvested by centrifugation at 700 × *g* for 3 minutes at 25°C and resuspended in fresh culture medium. For nucleic acid staining, cells were stained with 1-mM SYTO16 or 1-mM SYTO9 (Thermo Fisher Scientific) for 30 minutes in the dark, then washed and resuspended in fresh medium prior to imaging. After staining, the cells were washed again and resuspended in fresh culture medium for imaging. To observe the free locomotion of *C. reinhardtii*, two coverslips (0.13–0.17-mm thick, Matsunami) as spacers were placed on a microscope slide, and 5 mL of the culture containing densely growing cells was introduced between the spacers. A PTFE-PFA film (25-μm thick, NR5100-001, Flon Chemical) was placed on top to create a stable observation chamber. To observe the flowing *C. reinhardtii* cells, a cell culture was directly placed between a microscope slide and the objective lens. The flow was created by repeatedly tapping the culture with the side of a pipette during image recording.

#### Detection and tracking of swimming cells

Cell positions were detected and tracked in the autofluorescent chloroplast images using the real-time cell tracking algorithm with a minor modification. In this modification, segmentation was performed on the chloroplast images using OpenCV’s cv2.watershed() function. If the brightness-based tracking position fell within the segmented region, the centroid of the region was used to update the tracked position. Cell velocities were calculated from frame-to-frame displacements in the time series of cell positions acquired at a 1-kHz sampling rate. The position time series was smoothed using a Savitzky-Golay filter (window size, 50 ms; polynomial order, 1). The time series of cell movement speed was computed as a series of first derivatives of local polynomials fitted to the position data using the Savitzky–Golay filter. Cell orientations were calculated as angles between the *x*-axis and the projections onto the *xy* plane of vectors from the chloroplast centroids to the nuclear centroids. Prior to the vector calculation, the time series of the nuclear centroid positions was smoothed using a Savitzky-Golay filter (window size, 50 ms; polynomial order, 1), as was done for the chloroplast centroid positions.

### Strain list

*Caenorhabditis elegans* TOY1(*lite-1(ce314) gur-3(ok2245)* X;

Is[H20p::nls3::tdTomato,H20p::nls2::GCaMP6f])

*Hypsibius exemplaris* strain Z151

C*hlamydomonas reinhardtii* strain 137c

## Data and materials availability

Original volumetric image datasets for *C. reinhardtii* are available at https://doi.org/10.5281/zenodo.16919859. Other volumetric image datasets, which are not publicly available due to their large file sizes, are available from the corresponding author upon reasonable request. The codes used in this study for image acquisition and data analysis are available at https://github.com/hideharu-mikami-ISOP/ISOP-microscopy-LabVIEW-programs and https://github.com/MikamiLab/isop_microscopy_python_matlab. The real-time cell-tracking code is not publicly available in order to preserve the originality of a forthcoming publication, but will be provided by the corresponding author upon reasonable request.

## Acknowledgments

We thank M. Endo and A. Togashi for developing a prototype of an automated stage tracking system, M. Kusuzaki and M. Takei for fabricating a custom-built sample holder, R. Kamiya and T. Kato-Minoura for providing the *C. reinhardtii* strain, R. Okuyama and S. Maitani for their assistance in editing the Figures and Supplementary Videos, and Serendipity Lab for facilitating collaboration. The *C. vulgaris* used to feed the tardigrades was provided by courtesy of Chlorella Industry. This work was supported by JST PRESTO (JPMJPR1788), JST CREST (JPMJCR22N4, JPMJCR1653, JPMJCR12W1), JST FOREST (JPMJFR2248), JSPS KAKENHI (JP22H04926, JP22K19261, J19H03326, JP21H05279, JP21H05310, JP21K15139), The Okawa Foundation for Information and Telecommunications, Research Foundation for Opto-Science and Technology, The Uehara Memorial Foundation, The Asahi Glass Foundation, NTT-Kyushu University Collaborative Research, Joint Research by Exploratory Research Center on Life and Living Systems (ExCELLS program Nos. 19-501 and 22EXC601), and partly by research funds from the Yamagata Prefectural Government and Tsuruoka City, Japan.

## Author Contributions

H.M. conceived the idea, designed, built, and validated the ISOP microscope system, and obtained funding for the work. Y.Tomina, H.S., K.M., and H.M. carried out the imaging experiments. Y.Tomina, A.I., H.S., K.M., C.W., and H.M. analyzed the volumetric image data and stage tracking data. A.I., H.S., and R.H. developed the real-time cell tracking program. H.S. developed the automated stage tracking system. Y.Tomina, Y.Toyoshima, Y.I., and H.M. designed the experiments using *C. elegans*. Y.Toyoshima, M.K., K.K., Y.M., and S.Oe prepared *C. elegans* samples. Y.Toyoshima carried out imaging experiments using the spinning-disk confocal microscope and analyzed the neuronal signals and behaviors of *C. elegans* using TDE-RICA. Y.Tomina, A.I., Y.Toyoshima, Y.I., and H.M. interpreted the *C. elegans* data. Y.Tomina and S.T. prepared *H. exemplaris* samples. Y.Tomina, S.T., K.A., and H.M. designed the experiments using *H. exemplaris*. Y.Tomina, A.I., Y.Toyoshima, S.T., K.A., Y.I., and H.M. interpreted *H. exemplaris* data. Y.Tomina and H.M. designed the experiments using *C. reinhardtii*. Y.Tomina, Y.Y., and Y.N. prepared *C. reinhardtii* samples. Y.Tomina, A.I., Y.N., T.N., and H.M. interpreted the *C. reinhardtii* data. K.G. supervised optics-related work. K.I. and T.N. supervised *C. reinhardtii*-related work. K.A. supervised *H. exemplaris*-related work. S.Onami supervised image analysis-related work. Y.Toyoshima, T.I., and Y.I. supervised *C. elegans*-related work. Y.Tomina, A.I., and H.M. wrote the paper with assistance from all authors.

## References

1. Y. M. Sigal, R. Zhou, X. Zhuang, Visualizing and discovering cellular structures with super-resolution microscopy. Science 361, 880–887 (2018).

2. F. Randi, A. K. Sharma, S. Dvali, A. M. Leifer, Neural signal propagation atlas of *Caenorhabditis elegans*. Nature 623, 406–414 (2023).

3. Y. Wan et al., Single-Cell Reconstruction of Emerging Population Activity in an Entire Developing Circuit. Cell 179, 355–372.e23 (2019).

4. F. Pampaloni, E. G. Reynaud, E. H. K. Stelzer, The third dimension bridges the gap between cell culture and live tissue. Nat. Rev. Mol. Cell Biol. 8, 839–845 (2007).

5. T. L. Liu et al., Observing the cell in its native state: Imaging subcellular dynamics in multicellular organisms. Science 360 (2018).

6. Y. Wu et al., Multiview confocal super-resolution microscopy. Nature 600, 279–284 (2021).

7. J. Huisken, J. Swoger, F. Del Bene, J. Wittbrodt, E. H. K. Stelzer, Optical sectioning deep inside live embryos by selective plane illumination microscopy. Science 305, 1007–1009 (2004).

8. E. H. K. Stelzer et al., Light sheet fluorescence microscopy. Nat. Rev. Methods Primers 1, 1–25 (2021).

9. P. J. Keller, A. D. Schmidt, J. Wittbrodt, E. H. K. Stelzer, Reconstruction of zebrafish early embryonic development by scanned light sheet microscopy. Science 322, 1065–1069 (2008).

10. L. Sacconi, L. Silvestri, E. C. Rodríguez, G. A. B. Armstrong, F. S. Pavone, A. Shrier, G. Bub, KHz-rate volumetric voltage imaging of the whole Zebrafish heart. Biophys. Rep. 2, 100046 (2022).

11. E. H. Miyasaki, et al., High-speed 3D imaging with a 25-camera multifocus microscope. Optica 12, 1230–1241 (2025).

12. V. Voleti et al., Real-time volumetric microscopy of in vivo dynamics and large-scale samples with SCAPE 2.0. Nat. Methods 16, 1054–1062 (2019).

13. R. Prevedel et al., Simultaneous whole-animal 3D imaging of neuronal activity using light-field microscopy. Nat. Methods 11, 727–730 (2014).

14. Z. Lu et al., Long-term intravital subcellular imaging with confocal scanning light-field microscopy. Nat. Biotechnol. 43, 569–580 (2025).

15. Z. Lu et al., Virtual-scanning light-field microscopy for robust snapshot high-resolution volumetric imaging. Nat. Methods 20, 735–746 (2023).

16. Z. Wang et al., Kilohertz volumetric imaging of in-vivo dynamics using squeezed light field microscopy. Nat. Methods 22, 2194–2204 (2025).

17. B. J. Chang et al., Real-time multi-angle projection imaging of biological dynamics. Nat. Methods 18, 829–834 (2021).

18. J. Demas et al., High-speed, cortex-wide volumetric recording of neuroactivity at cellular resolution using light beads microscopy. Nat. Methods 18, 1103–1111 (2021).

19. V. Venkatachalam et al., Pan-neuronal imaging in roaming *Caenorhabditis elegans*. Proc. Natl. Acad. Sci. U.S.A. 113, E1082–E1088 (2016).

20. J. P. Nguyen et al., Whole-brain calcium imaging with cellular resolution in freely behaving *Caenorhabditis elegans*. Proc. Natl. Acad. Sci. U.S.A. 113, E1074–E1081 (2016).

21. A. Atanas et al., Brain-wide representations of behavior spanning multiple timescales and states in *C. elegans*. Cell 186, 4134–4151.e31 (2023).

22. C. Dunsby, Optically sectioned imaging by oblique plane microscopy. Opt. Express 16, 20306 (2008).

23. M. Kumar, S. Kishore, J. Nasenbeny, D. L. McLean, Y. Kozorovitskiy, Integrated one- and two-photon scanned oblique plane illumination (SOPi) microscopy for rapid volumetric imaging. Opt. Express 26, 13027 (2018).

24. B. Yang et al., DaXi—high-resolution, large imaging volume and multi-view single-objective light-sheet microscopy. Nat. Methods 19, 461–469 (2022).

25. C. J. R. Sheppard, Super-resolution in confocal imaging. Optik (Stuttgart) 80, 53–54 (1988).

26. Y. Gong, Y. Tian, C. Baker, A fully water coupled oblique light-sheet microscope. Sci. Rep. 12, 1–11 (2022).

27. T. Schrödel, R. Prevedel, K. Aumayr, M. Zimmer, A. Vaziri, Brain-wide 3D imaging of neuronal activity in *Caenorhabditis elegans* with sculpted light. Nat. Methods 10, 1013–1020 (2013).

28. Y. Toyoshima et al., Ensemble dynamics and information flow deduction from whole-brain imaging data. PLoS Comput. Biol. 20, e1011848 (2024).

29. V. Susoy et al., Natural sensory context drives diverse brain-wide activity during *C. elegans* mating. Cell 184, 5122–5137.e17 (2021).

30. C. Wen et al., 3DeeCellTracker, a deep learning-based pipeline for segmenting and tracking cells in 3D time lapse images. eLife 10, e59187 (2021).

31. J. A. Nirody, L. A. Duran, D. Johnston, D. J. Cohen, Tardigrades exhibit robust interlimb coordination across walking speeds and terrains. Proc. Natl. Acad. Sci. U.S.A. 118, e2107289118 (2021).

32. M. R. Smith, J. Ortega-Hernández, *Hallucigenia*’s onychophoran-like claws and the case for Tactopoda. Nature 514, 363–366 (2014).

33. S. Tanaka, K. Aoki, K. Arakawa, In vivo expression vector derived from anhydrobiotic tardigrade genome enables live imaging in Eutardigrada. Proc. Natl. Acad. Sci. U.S.A. 120, e2216739120 (2023).

34. V. Gross, G. Mayer, Cellular morphology of leg musculature in the water bear Hypsibius exemplaris (Tardigrada) unravels serial homologies. R. Soc. Open Sci. 6, 191159 (2019).

35. K. Drescher, K. C. Leptos, R. E. Goldstein, How to track protists in three dimensions. Rev. Sci. Instrum. 80, 014301 (2009).

36. M. Polin, I. Tuval, K. Drescher, J. P. Gollub, R. E. Goldstein, *Chlamydomonas* swims with two “gears” in a eukaryotic version of run-and-tumble locomotion. Science 325, 487–490 (2009).

37. J. B. Raina, V. Fernandez, B. Lambert, R. Stocker, J. R. Seymour, The role of microbial motility and chemotaxis in symbiosis. Nat. Rev. Microbiol. 17, 284–294 (2019).

38. S. Dupuis, S. S. Merchant, *Chlamydomonas reinhardtii*: a model for photosynthesis and so much more. Nat. Methods 20, 1441–1442 (2023).

39. E. M. Purcell, Life at low Reynolds number. Am. J. Phys. 45, 3–11 (1977).

40. T. W. Su, L. Xue, A. Ozcan, High-throughput lensfree 3D tracking of human sperms reveals rare statistics of helical trajectories. Proc. Natl. Acad. Sci. U.S.A. 109, 16018–16022 (2012).

41. J. Alvelid, M. Damenti, C. Sgattoni, I. Testa, Event-triggered STED imaging. Nat. Methods 19, 1268–1275 (2022).

42. D. Mahecic et al., Event-driven acquisition for content-enriched microscopy. Nat. Methods 19, 1262–1267 (2022).

43. Y. Shi et al., Smart lattice light-sheet microscopy for imaging rare and complex cellular events. Nat. Methods 21, 301–310 (2024).

44. Z. Zhang, L. E. Russell, A. M. Packer, O. M. Gauld, M. Häusser, Closed-loop all-optical interrogation of neural circuits in vivo. Nat. Methods 15, 1037–1040 (2018).

45. N. Friedman, A. Kaplan, N. Davidson, Acousto-optic scanning system with very fast nonlinear scans. Opt. Lett. 25, 1762–1764 (2000).

46. V. R. Prasanna, A. S. Mutlu, C. W. Meng, Label-free biomedical imaging of lipids by stimulated Raman scattering microscopy. Curr. Protoc. Mol. Biol. 109, 30.3.1–30.3.17 (2015).

47. C. Wen, Deep Learning-Based Cell Tracking in Deforming Organs and Moving Animals. Methods Mol. Biol. 2800, 203–215 (2024).

48. T. Ahamed, A. C. Costa, G. J. Stephens, Capturing the continuous complexity of behaviour in *Caenorhabditis elegans*. Nat. Phys. 17, 275–283 (2021).

49. A. Matsumoto et al., Neuronal sensorimotor integration guiding salt concentration navigation in *Caenorhabditis elegans*. Proc. Natl. Acad. Sci. U.S.A. 121, e2310735121 (2024).

50. D. B. Stern, The Chlamydomonas Sourcebook: Organellar and Metabolic Processes (Academic Press, Oxford, ed. 2, 2008).

