## Supplementary Information for "Image-scanning light-sheet microscopy for high-speed volumetric imaging of complex biological dynamics"

**Supplementary Note S1: Comparison of imaging sensitivity between image-scanning selective plane illumination microscopy and image-scanning digital scanned light-sheet microscopy**

The key difference between selective plane illumination microscopy (SPIM) and digital scanned light-sheet microscopy (DSLM)—two principal implementations of light-sheet microscopy—lies in the spatiotemporal profile of the excitation beam, which leads to a substantial difference in imaging sensitivity when implementing image-scanning–based multi-plane imaging. We quantitatively evaluate this difference using a simplified model.

Let the field of view along the $y$ direction (the direction perpendicular to the excitation beam propagation direction in the focal plane) be $l$. In image-scanning DSLM, the excitation beam is assumed to have uniform intensity with a width $d$ along the $y$ direction and to be scanned in the same direction. In image-scanning SPIM, by contrast, the excitation light sheet is assumed to have uniform intensity with a width equal to the entire field of view, i.e., $l$, in the $y$ direction. In image-scanning SPIM, the maximum permissible stroboscopic illumination time without noticeable motion blur is typically taken as the time for the optical image to move by one pixel, or more rigorously, a distance smaller than the size of point spread function (PSF). In image-scanning DSLM, to avoid motion blur, the excitation beam must traverse the corresponding pixel within the time the optical image shifts by one pixel. In both cases, the excitation duration for a fluorophore is the same—namely, the time for the optical image to move by one pixel. However, assuming equal power, the excitation beam in image-scanning DSLM has an energy density higher than that in image-scanning SPIM by a factor of $l/d$. Therefore, the imaging sensitivity in image-scanning DSLM is higher by a factor equal to the ratio of the field of view to the beam width along the $y$ direction. In the design of our DSLM-based image-scanning oblique plane (ISOP) microscope, the Gaussian beam waist radius ($1/e^2$ intensity) along the $y$ direction was ~3 µm, corresponding to $d$ = 6 µm, while the field of view in the $y$ direction $l$ ranged from 50 µm to 287 µm in the imaging experiments, yielding an estimated sensitivity enhancement of 8.4–48-fold. This indicates that the required excitation laser power differs by more than an order of magnitude between image-scanning SPIM and image-scanning DSLM. As a result, the DSLM-based ISOP microscope achieved sufficient imaging sensitivity with laser output power below 100 mW, which is readily available from commonly used laser sources in fluorescence microscopes. In other scanning configurations (Supplementary Note S6), the effect of motion blur must be carefully considered by taking into account the magnification change in the image-scanning direction, but the theoretical PSF evaluation results (Supplementary Note S2) indicate comparable PSF degradations across configurations, suggesting that the above discussion is applicable irrespective of the scanning configuration under practical implementations.

**Supplementary Note S2: Evaluation of the spatial resolution**

We evaluated the spatial resolution of the ISOP microscope using fluorescent beads and theoretical calculations. 200-nm fluorescent beads (Fluoresbrite YG Microspheres – 0.20 µm, Polysciences, Inc.) were embedded in agarose gel and sealed with a 25-µm-thick perfluoroalkoxy (PFA) film. Imaging was performed at a magnification of 31.7 using a 200-mm-focal-length tube lens (TTL200-A, Thorlabs), corresponding to a pixel size of 205 nm. For each bead, line profiles were extracted along the $x$, $y$, and $z$ axes, centered at the voxel with maximum intensity. These profiles were fitted with an offset Gaussian function, and the full width at half maximum (FWHM) in each direction was calculated from the fits.

In addition to assessing the dependence of the PSF size on the scanning velocity ratio ($\rho$), we also evaluated its dependence on the $z$ position to verify the performance of the remote focusing system (Fig. S20). Fluorescent bead images were acquired over a $z$ range of –50 µm to +50 µm by moving the sample, and FWHM values were measured along the $x$, $y$, and $z$ directions, as well as along the theoretical PSF axes $X$ and $Z$ in the $xz$ plane, which was inclined at 23°.

Theoretical PSFs were derived using scalar diffraction theory with the Debye approximation, incorporating the designed excitation beam width and the solid angle of fluorescence collection. The corresponding FWHM values were calculated for the $x$, $y$, and $z$ directions for comparison. In these experiments, the scanning velocity ratio was set to 12.77. As shown in Fig. S20, the measured FWHM values were slightly larger than the theoretical predictions in the *x*, *y*, and *z* directions. Nonetheless, uniform spatial resolution was maintained across the full *z* range of –50 μm to +50 μm. Submicrometer lateral resolution was achieved. Along the *Z* axis, the FWHM value was 2,136 ± 204 nm (mean ± s.d., *n* = 170 beads), representing the direction with the lowest resolution.

Furthermore, theoretical PSFs were calculated for a range of scanning velocity ratios and imaging configurations. Fig. S21*A* shows PSF profiles in the *xy* and *xz* planes for various values of the scanning velocity ratio ($\rho$). As $\rho$ decreased, the profiles broadened in the *x* and *y* directions, which corresponded to increased motion blur, and the inclination angle of the PSF in the *xz* plane changed accordingly. Theoretical FWHM in the *x*, *y*, and *z* directions for different $\rho$ values are presented in Fig. S21*B*, exhibiting trends consistent with the experimental results shown in Fig. 1*F*. Fig. S21*C* shows the theoretical FWHM values for alternative image scanning configurations depicted in Fig. S22 (*A* and *B*) (see Supplementary Note S6). In these configurations, $\rho > 0$ denotes alignment of the image scanning direction with the beam scanning direction (Fig. S22*B*), while $\rho < 0$ indicates the opposite alignment (Fig. S22*A*). Decreasing the absolute value of $\rho$ significantly increased the FWHM in the *y* direction. Although the spatial resolution in the *y* direction can be improved under certain conditions, as in conventional image scanning microscopy (*1, 2*), such improvement was not observed in the ISOP microscope setup due to the need for an excitation beam with a relatively large width to suppress divergence along the beam propagation axis.

**Supplementary Note S3: Motion-induced image deformation**
Imaging modalities such as light-sheet and confocal microscopy, which sequentially acquire voxel values within a volume, are subject to image deformation if the object moves during single volumetric image acquisition. This deformation can be negligible when the object's movement is slow, but becomes nontrivial at higher velocities, necessitating consideration of both the type and magnitude of deformation.

For simplicity, we assume infinitely fast beam scanning and a constant object movement speed, denoted as $v_{\mathrm{o}}$. When the object's motion is oriented perpendicular to the focal plane scanning direction, shear distortion occurs (Fig. S23*A*). When the motion is oriented parallel to the focal plane scanning direction, either expansion or shrinkage is observed (Fig. S23, *B* and *C*). In general, a combination of these deformations arises depending on the motion direction. The deformation magnitude is determined by the ratio of the object movement speed to the focal plane scanning speed $v_{\mathrm{s}}$. For example, in the case of expansion or shrinkage, the width of the object along the motion direction is scaled by a factor of $\frac{1}{1-\frac{v_{\mathrm{o}}}{v_{\mathrm{s}}}}$, with the sign of $v_{\mathrm{o}}$ defined relative to the direction of focal plane scanning. For shear deformation, the apparent length increases by a factor of $\sqrt{1+\left(\frac{v_{\mathrm{o}}}{v_{\mathrm{s}}}\right)^2}$. Since $v_{\mathrm{s}}$ is a controllable microscope parameter, and $v_{\mathrm{o}}$ can be estimated from inter-volume displacement of the object divided by the volume period, the amount of deformation can be estimated from inter-volume displacement between consecutive volumetric images. Although the focal plane scanning speed in ISOP microscopy is not necessarily constant and includes backward motion during frame acquisition, the above formulation still provides a practical approximation. This approximation model is also applicable when the focal plane scanning direction is not perpendicular to the focal plane, as is the case in ISOP microscopy. In the *H. exemplaris* imaging experiment, the maximum estimated movement speed of muscle fibers and the focal plane scanning speed were ~300 μm $s^{-1}$ (predominantly horizontal) and ~3 mm $s^{-1}$, respectively. Based on the deformation model described above, these values correspond to the errors in the muscle fiber length of $1/(1 \pm 300/3000) - 1 \sim \pm 10\%$ when the motion was parallel to the focal plane scanning direction, and $\sqrt{1+0.1^2} - 1 \sim 0.5\%$ when the motion was perpendicular to it.

Moreover, in cases such as imaging of *Caenorhabditis elegans* neurons and *Chlamydomonas reinhardtii* cells, where the object (i.e., an individual cell) is fully contained within a local volumetric region that corresponds to a single image frame, the motion-induced deformation is further reduced. This is because the effective scanning speed during frame acquisition is higher due to the absence of temporal gaps between adjacent image frames. For example, in our *C. elegans* imaging, assuming a cell movement speed of 300 μm $s^{-1}$ and a focal plane scanning speed of 23 mm $s^{-1}$, the estimated distortion was within ±1.3%. For *C. reinhardtii* imaging, with a locomotion speed of 10 mm $s^{-1}$ and a focal plane scanning speed of 200 mm $s^{-1}$, the estimated distortion was within ±5%. However, when an object spans multiple adjacent local volumetric regions that correspond to different image frames, discontinuous image displacement can occur at their boundaries due to relatively long exposure time intervals (Fig. S23*D*). In our *C. elegans* imaging, the estimated displacement was ~0.48 μm for an object moving at 300 μm $s^{-1}$. This displacement was considered negligible, as it is much smaller than the typical nucleus diameter (~3 μm). In the case of *C. reinhardtii* imaging, where a full volumetric image is reconstructed from a single image frame, such discontinuous deformation did not occur.

**Supplementary Note S4: Quasi-continuous sheet-shaped beam generation**
In conventional selective plane illumination microscopy, a sheet-shaped excitation beam is typically generated by beam shaping with cylindrical lenses. To replicate this using an acousto-optic deflector (AOD), the simplest approach involves restricting the effective aperture of the AOD along the acoustic wave propagation axis (i.e., the extent of an isolated sound wave). However, this method yields extremely low diffraction efficiency, rendering it impractical for most applications.

A more viable strategy for generating a pseudo-sheet-shaped beam while maintaining high net diffraction efficiency is to rapidly scan an array of beam spots. Techniques for producing such arrays using an AOD have been demonstrated in previous studies (*3, 4*). This can be accomplished by driving the AOD with a multi-tone signal composed of evenly spaced frequency components with quadratic phase variation. By periodically shifting the entire array along its axis by an amount equal to the inter-spot spacing, a quasi-continuous sheet-shaped beam can be synthesized. This is achieved by uniformly modifying the frequencies of the tone components in the multi-tone signal. In configurations using two AODs, a pseudo-sheet-shaped beam can be generated by driving one AOD with the multi-tone signal and the other with a monotone signal.

Leveraging the programmability of AOD modulation, the system can seamlessly switch between image-scanning LSM, selective plane illumination microscopy, and projection imaging. Notably, this transition is accomplished solely through changes to the driving signal, without any mechanical adjustments or alterations to the optical path.

**Supplementary Note S5: Optical system design**
To evaluate the imaging performance of the designed optical system, we performed ray-tracing simulations for each relay optical system in the optical setup using OpticStudio, as shown in Fig. S24. All the relay systems exhibited rms wavefront aberrations well below the diffraction limit (0.07 λrms) over practical field angles (0 to 0.033 degrees for excitation beam from AOD2 and 0 to 1 degree for fluorescence light paths). Although the simulation was performed using the full aperture of the objective lenses, a portion of the aperture was used for fluorescence image formation due to the oblique plane imaging configuration, indicating that the wavefront aberrations under the actual imaging conditions would be smaller than those simulated with the full aperture. We did not perform the numerical simulation of the entire optical system because no lens model was available for the objective lenses, but the above results support the validity of the optical design.

**Supplementary Note S6: Alternative scanning configurations**
Alternative configurations in which the beam scanning direction is aligned with or opposed to the image scanning direction are illustrated in Fig. S22. The sectioned images are not skewed but appear compressed or expanded along the scanning axis. These configurations result in slight

magnification variation along the image scanning direction, similar to conventional image scanning microscopy. Nevertheless, accurate volumetric reconstruction remains feasible by adjusting the scale of the sectioned images according to the specified magnification.

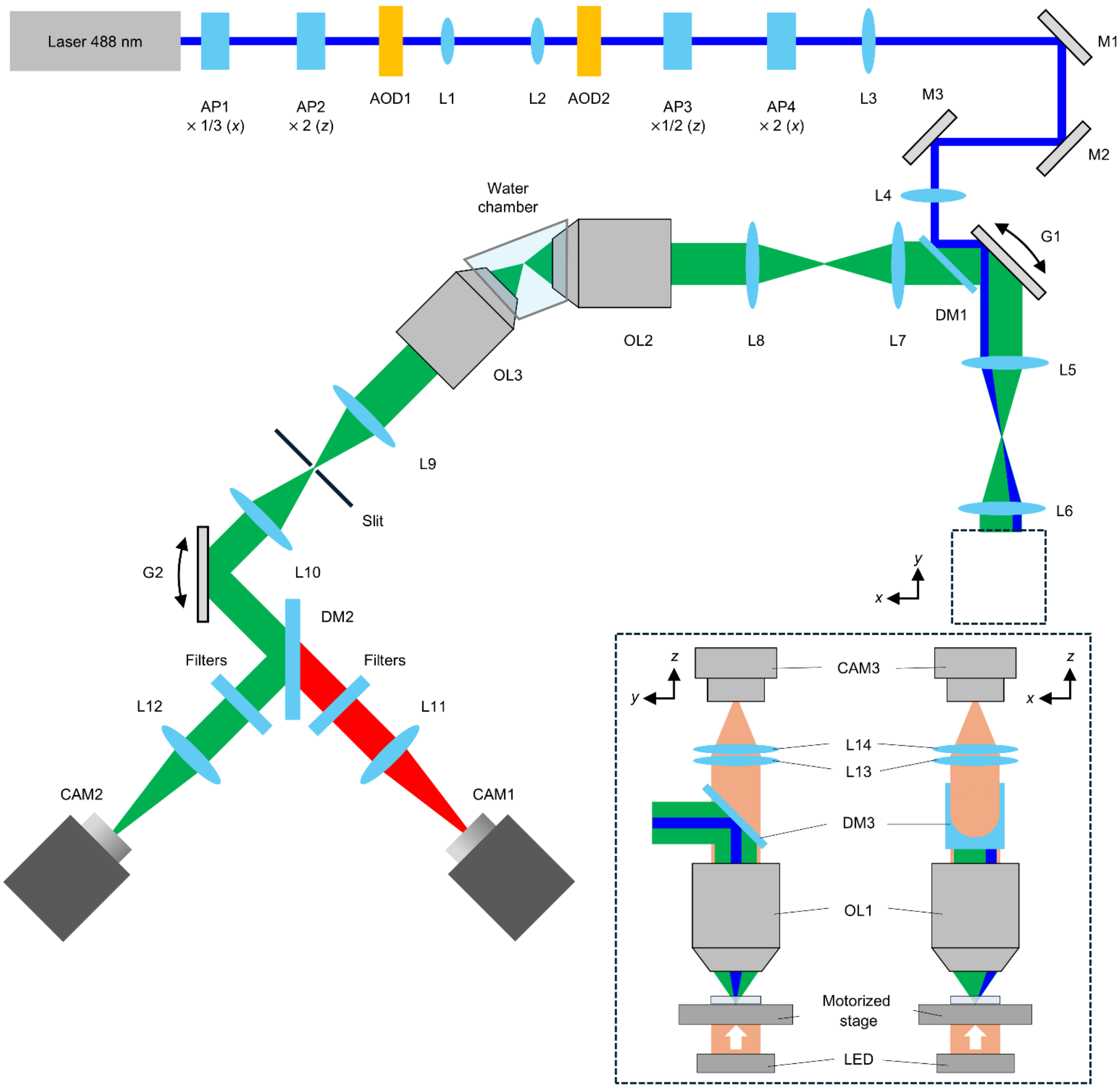


**Fig. S1. Detailed optical layout of the ISOP microscope.** AP1-AP4, anamorphic prisms. AOD1, AOD2, acousto-optic deflectors. L1-L14, lenses. DM1-DM3, dichroic mirrors. OL1-OL3, objective lenses. G1, G2, galvanometric scanners. CAM1-CAM3, cameras. The inset shows side views of the microscope around the sample.

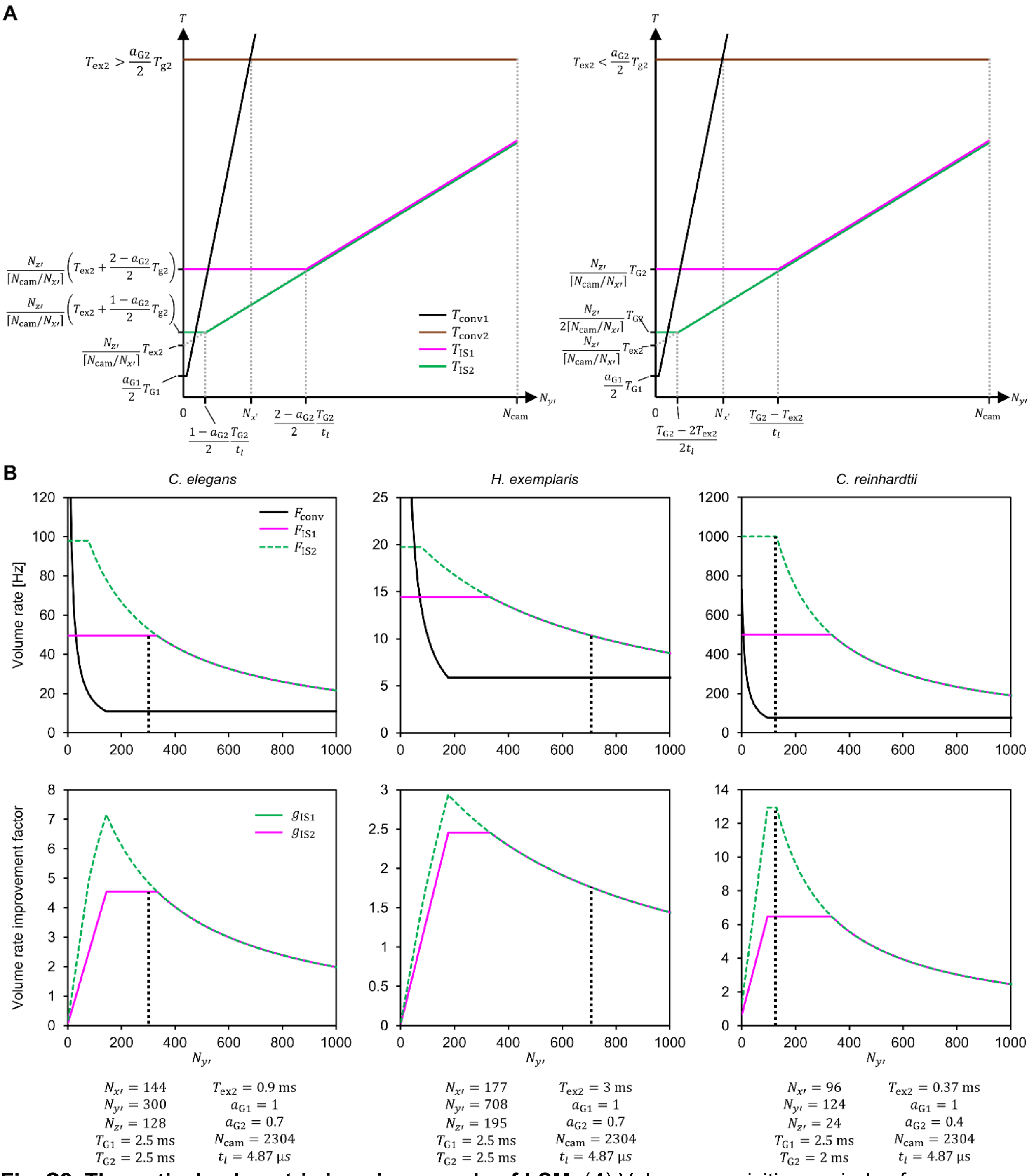


**Fig. S2. Theoretical volumetric imaging speeds of LSM.** (*A*) Volume acquisition periods of conventional and image-scanning LSM at different imaging configurations. (*B*) Volume rates, volume rate enhancement factors achieved through image scanning, and parameters used for imaging *C. elegans* (left), *H. exemplaris* (center), and *C. reinhardtii* cells (right). The vertical dotted lines indicate the values of $N_{y'}$ used. See Methods for the notation.

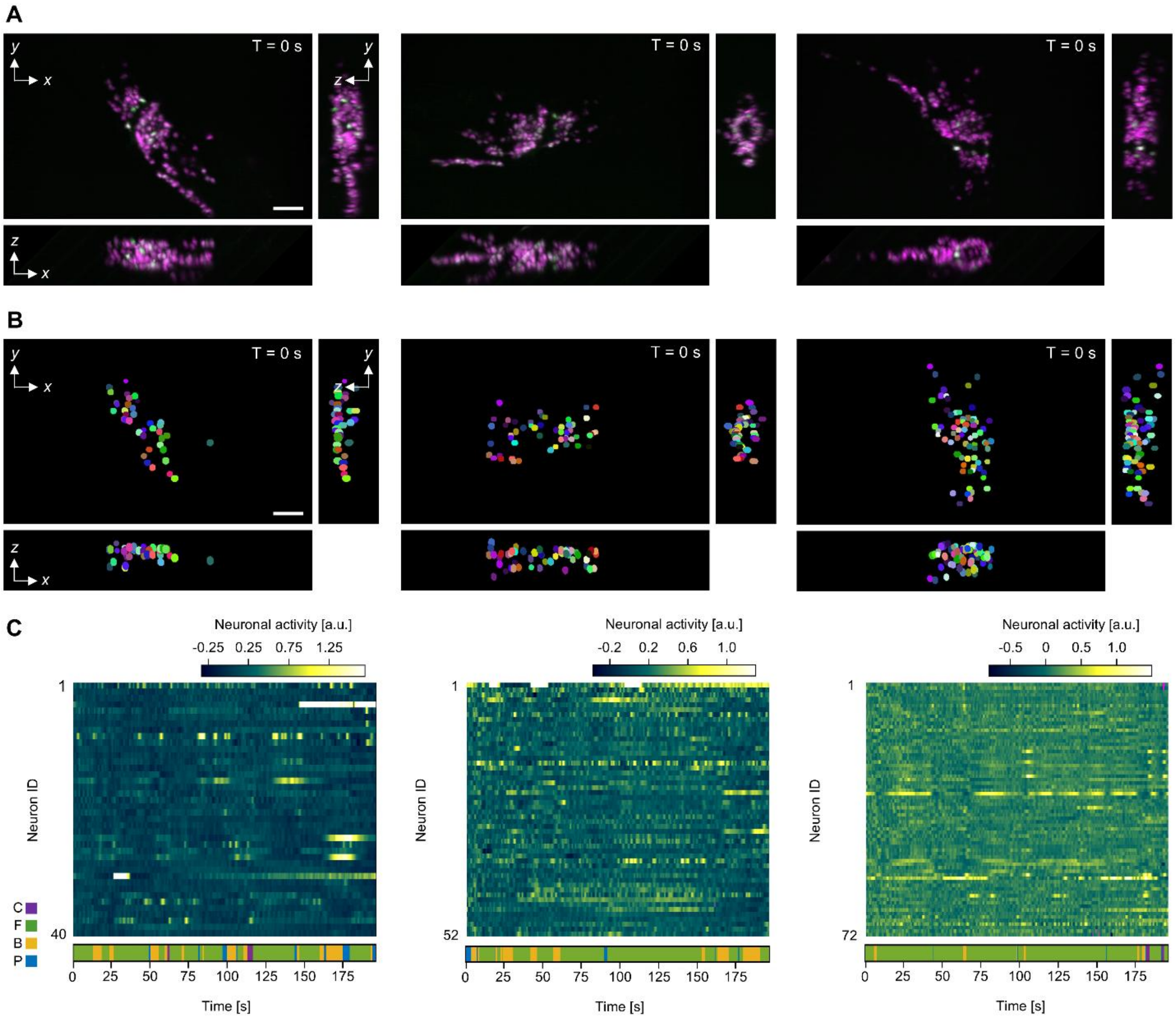


**Fig. S3. Whole-brain calcium imaging of freely behaving *C. elegans* using ISOP microscopy (Worms 2-4).** (*A*) Maximum intensity projections in the *xy*, *xz*, and *zy* planes of a volumetric image acquired at 50 vps from the head regions of freely behaving *C. elegans* (see Movie S3). Scale bar, 20 µm. (*B*) Cell detection results. Scale bar, 20 µm. (*C*) Time series of neuronal activity (top) and manually annotated behavioral states (bottom). All panels use the same format as Fig. 2 (*A* to *C*). The columns from left to right correspond to Worms 2, 3, and 4, respectively. A few neurons exceed the upper limit of the display color scale and therefore appear saturated in the activity map shown in (*C*) (Neurons 4 and 31 in Worm 2; Neuron 1 in Worm 3).

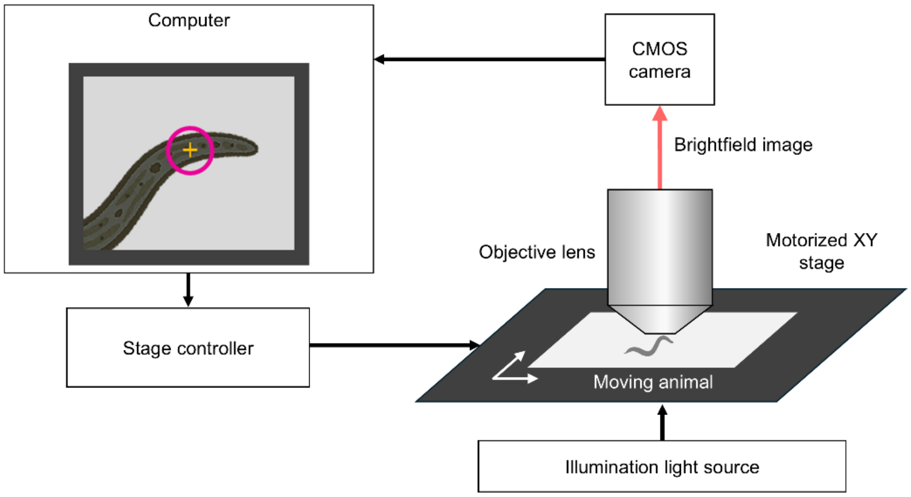


**Fig. S4. Automated stage tracking system.** Brightfield images of the specimen are analyzed in real time, and the resulting displacement is compensated by driving a motorized XY stage.

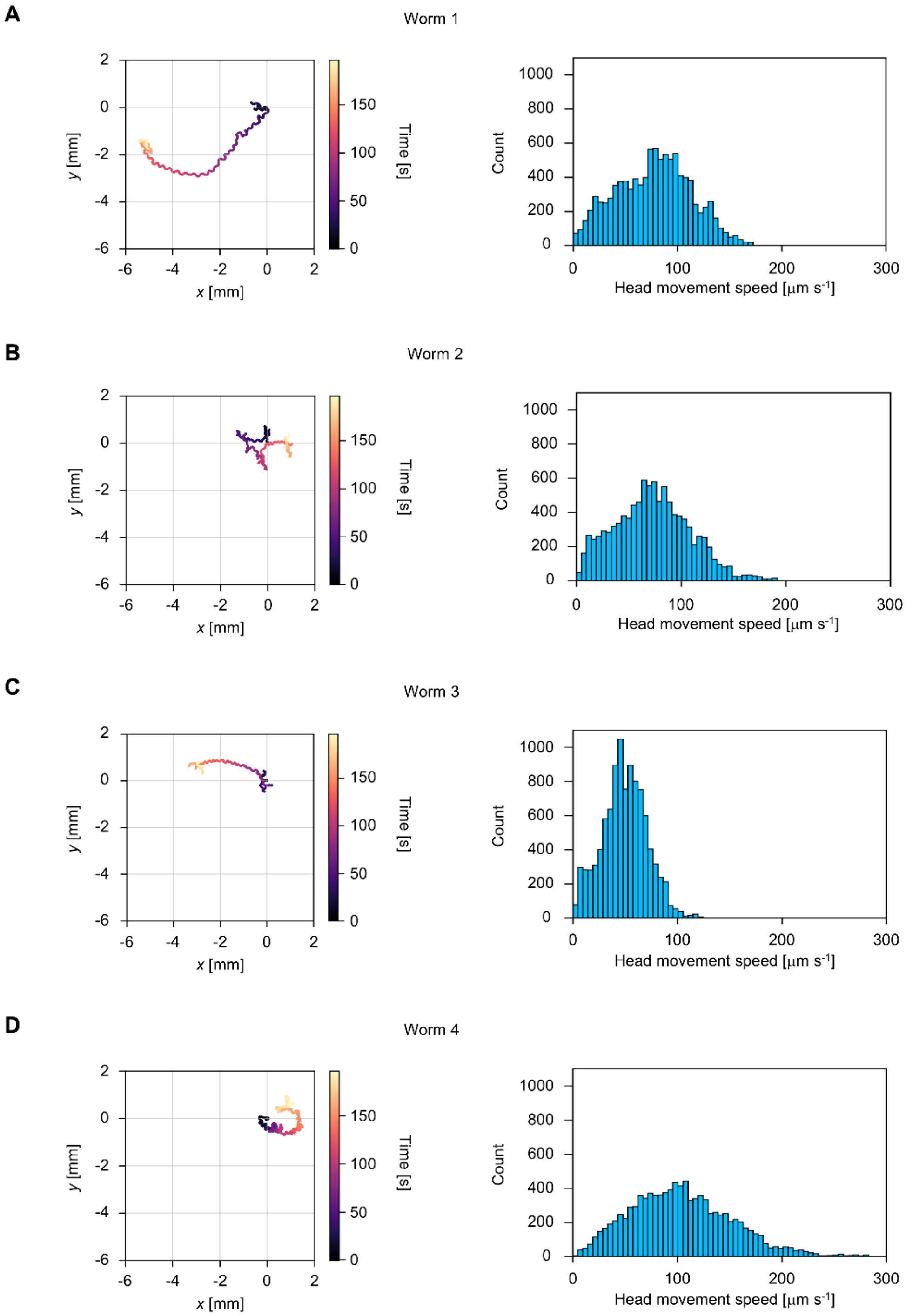


**Fig. S5. Tracking trajectories and head movement speed distributions of *C. elegans*.** Trajectories of the heads of *C. elegans* tracked over 196.2 s, color-coded by time (left), and histograms of head speed (right). Each trajectory was generated from smoothed position data recorded by the automated stage tracking system. The smoothing was performed using a Savitzky–Golay filter (window size, 1 s; polynomial order, 1). Each speed was computed as the magnitude of a velocity vector obtained as the first derivative of a local polynomial fitted to the position data using the Savitzky–Golay filter. (*A*)–(*D*) correspond to Worms 1–4, respectively.

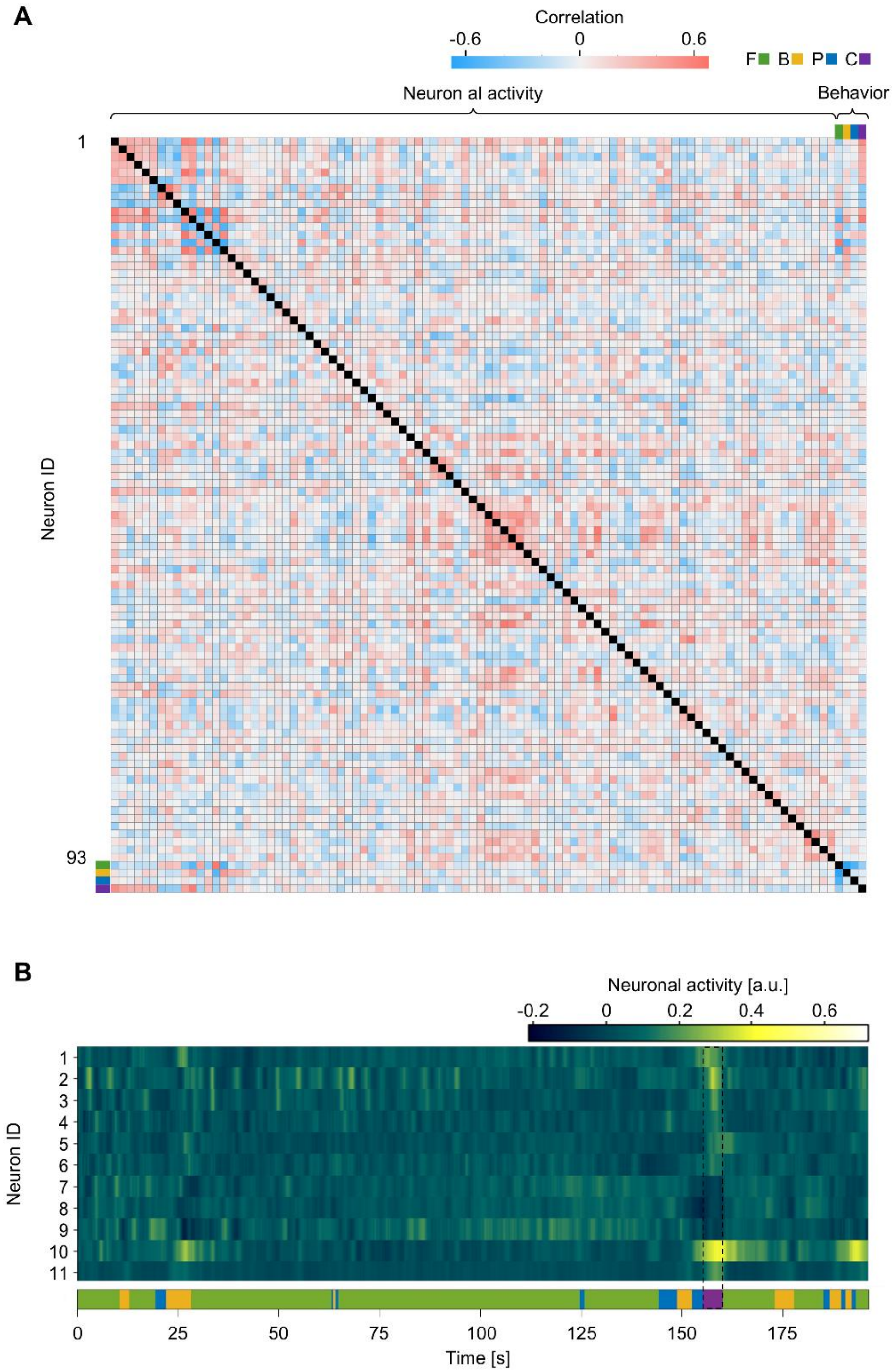


**Fig. S6. Correlation between neuronal activity and behavior in *C. elegans* (Worm 1).** (*A*) Heatmap showing pairwise Pearson correlation coefficients between 93 neurons with detected neuronal activity and behavioral states manually annotated from brightfield images in Worm 1, including forward locomotion (green), backward locomotion (yellow), pausing (blue), and coiling (purple). Correlation values are color-coded (red, positive; blue, negative). Diagonal elements (black) indicate self-correlation. (*B*) Time series of neuronal activity from 11 neurons correlated with coiling behavior ($|r|>0.3$), highlighting prominent activity level changes during coiling behavior (155.1–159.9 s).

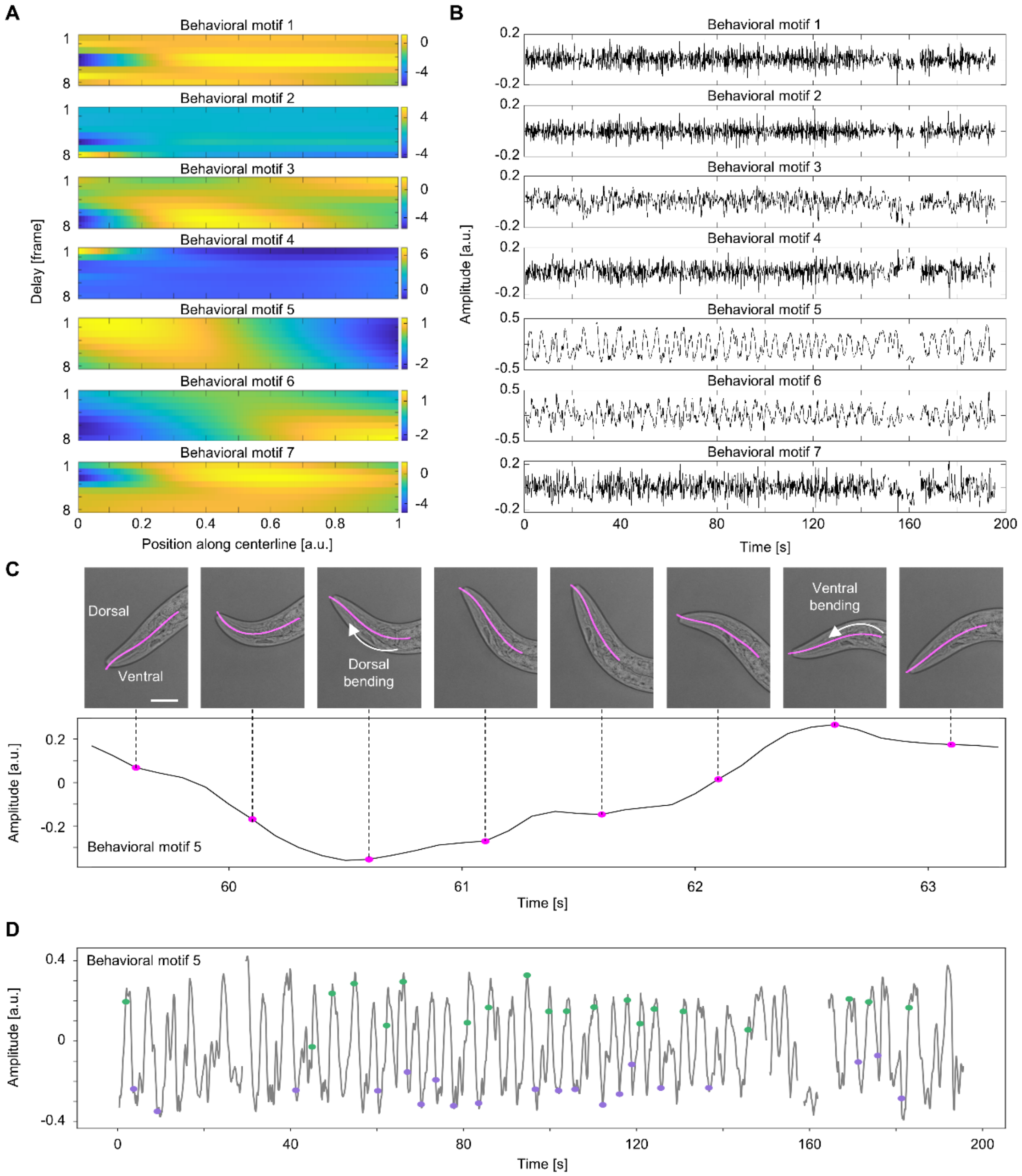


**Fig. S7. Quantitative analysis of head posture in *C. elegans*.**
(*A*) Seven behavioral motifs extracted from the time series of Worm 1's head posture. The color represents normalized angles. The horizontal and vertical axes show the position along the centerline from the nose tip (0) to the neck (1) and delay, respectively. (*B*) Time series of behavioral motif amplitudes (indicating temporal occurrence of each behavioral motif). (*C*) Time series of behavioral motif 5 amplitudes and corresponding head postures overlaid with the extracted centerline shown as magenta curves, illustrating the correlation between the worm's head bending direction and the amplitude (ventral and dorsal bending correspond to negative and positive values, respectively). Scale bar, 20 μm. (*D*) Time series of behavioral motif 5 amplitudes, with green and purple marks indicating time points at which maximal ventral and dorsal bending postures were identified by visual inspection of brightfield images, illustrating the correlation between the worm's head bending direction and the amplitude as in (*C*).

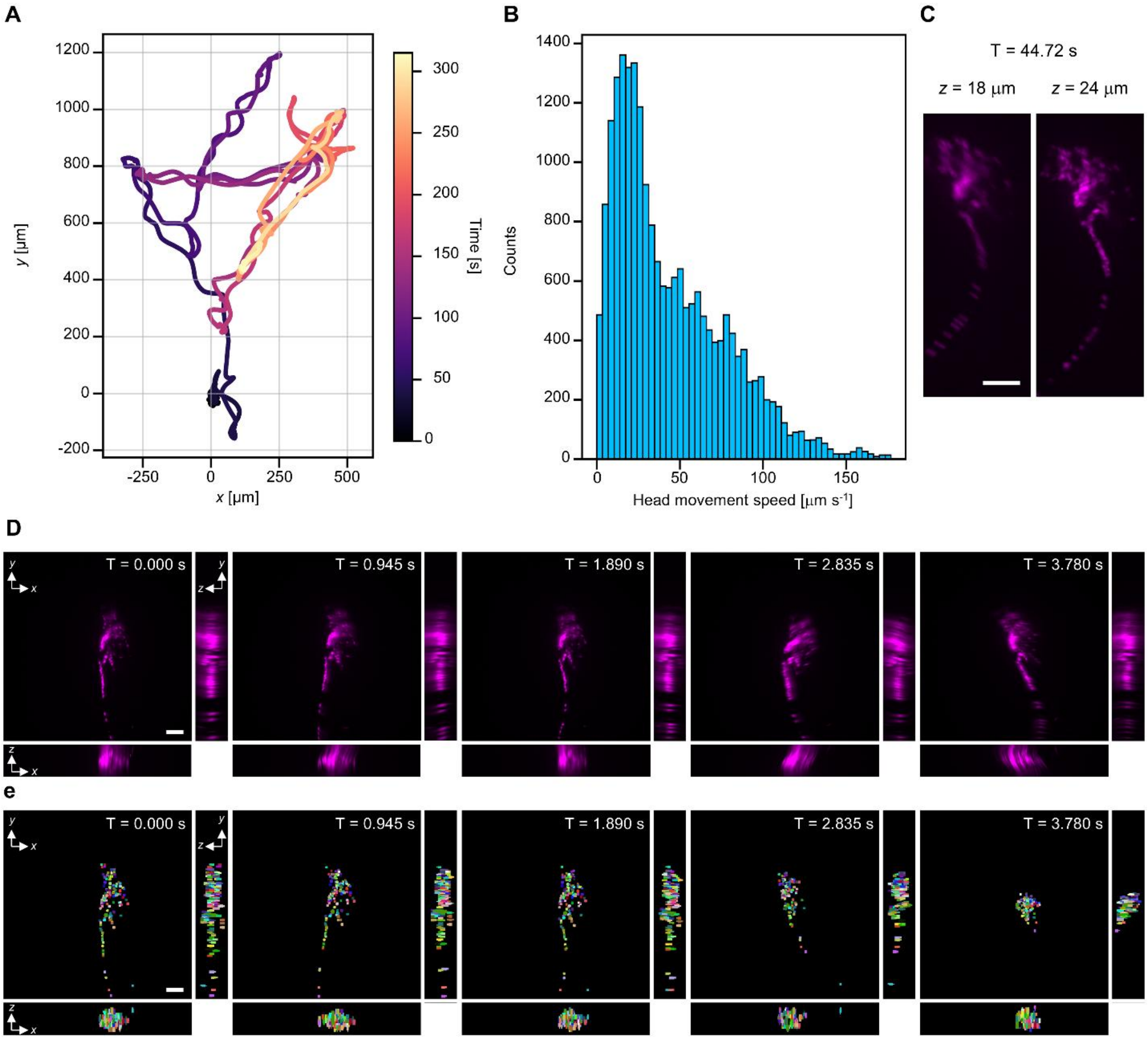


**Fig. S8. Whole-brain calcium imaging of a freely behaving *C. elegans* using spinning-disk confocal microscopy.** (*A*) Trajectory of the head of the worm analyzed. A time course from 0 to 315 s is represented by a color gradient. The trajectory was generated from smoothed position data recorded by the automated stage tracking system integrated with the spinning-disk confocal microscope. The smoothing was performed using a Savitzky–Golay filter (window size, 1 s; polynomial order, 1). (*B*) Histogram of the worm's head speed. Each speed was computed as the magnitude of a velocity vector obtained as the first derivative of a local polynomial fitted to the position data using the Savitzky–Golay filter (window size, 1 s; polynomial order, 1). (*C*) Slice image of the tdTomato fluorescence showing prominent motion blur (left, T = 44.72 s, *z* = 18 μm), clearly distinguishable when compared to another slice image at a different *z* position (right, T = 44.72 s, *z* = 24 μm). (*D*) Time series of Maximum intensity projections (MIPs) of the tdTomato fluorescence volumetric images, showing significant deformation artifacts and displacement of neurons. The images were resampled in the *z* direction with linear interpolation before MIP generation to better visualize motion-induced deformation. (*E*) Cell tracking results, showing significant tracking errors at T = 3.780 s. Scale bars: 20 μm.

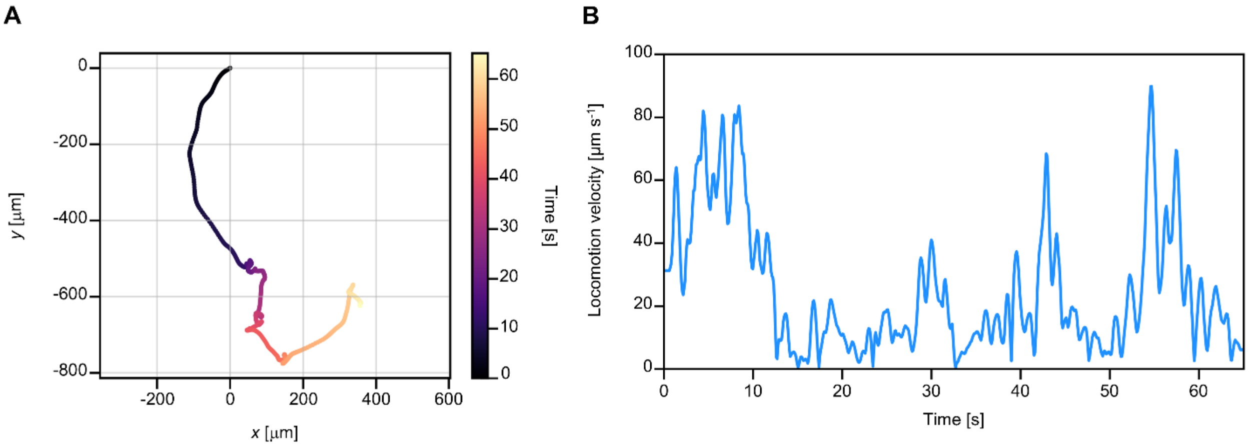


**Fig. S9. Locomotion trajectory and speed of the tardigrade.** (*A*) Locomotion trajectory of the tardigrade, recorded using the automated stage tracking system. The colors indicate elapsed time (0–64.8 s). The trajectory was generated from a position time series smoothed using a Savitzky-Golay filter (window size, 1 s; polynomial order, 1). (*B*) Time series of locomotion speed, computed as the magnitude of velocity vectors obtained as first derivatives of local polynomials fitted to the position data using the Savitzky–Golay filter.

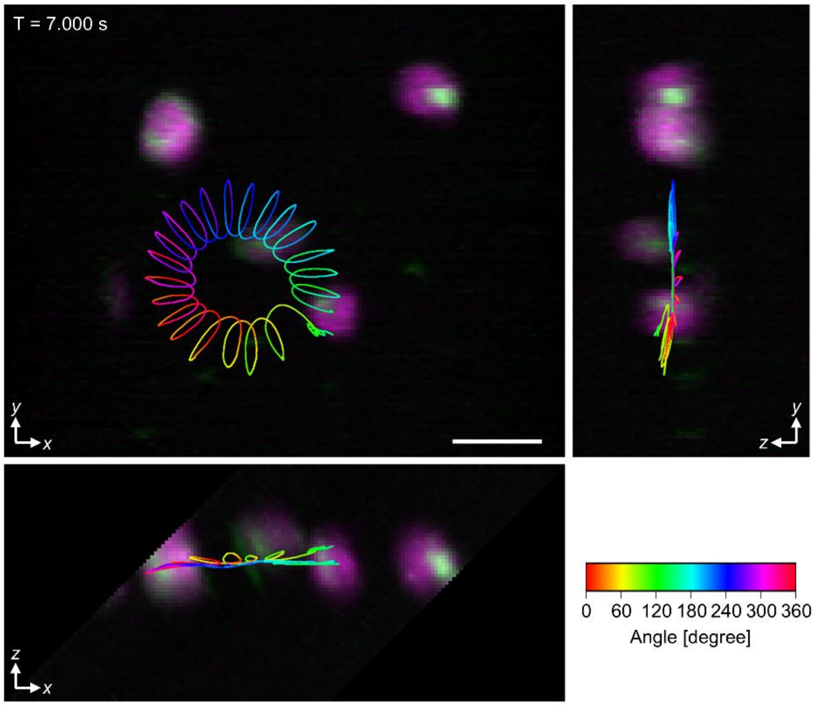


**Fig. S10. Orientations of a *C. reinhardtii* cell.** Maximum intensity projections in the *xy*, *xz*, and *zy* planes with the trajectory, both identical to those shown in Fig. 4*A*, are displayed except that the trajectory is color-coded by orientation in the *xy* plane. Scale bar, 10 µm.

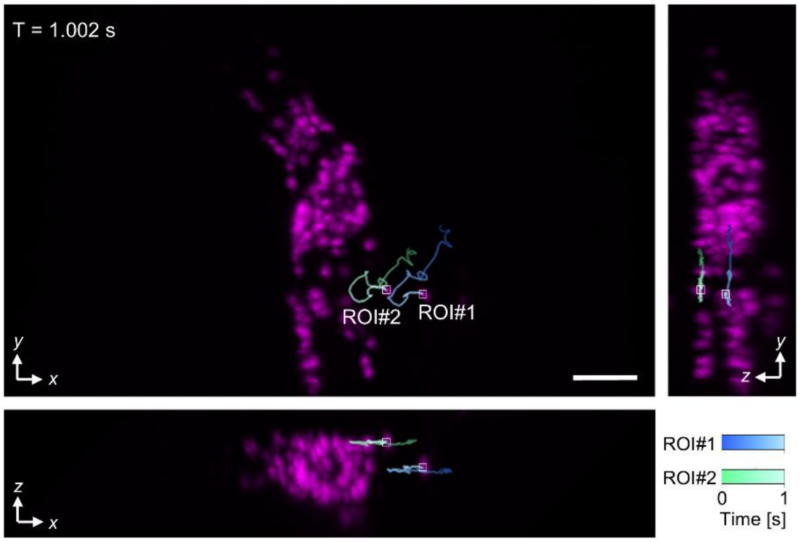


**Fig. S11. Real-time tracking of tdTomato-labeled neurons in freely behaving *C. elegans*.** Maximum intensity projections of a volumetric image in the *xy*, *xz*, and *zy* planes acquired at 50 vps from Worm 2, showing two tdTomato-labeled neurons tracked in real time. The trajectories of these cells over the preceding 1 s are shown in green and blue. The cells were continuously tracked for 33.84 s within a total recording session of 196.2 s, until tracking ceased due to tracking errors. See Movie S8 for the full tracking sequence. Scale bar, 20 µm.

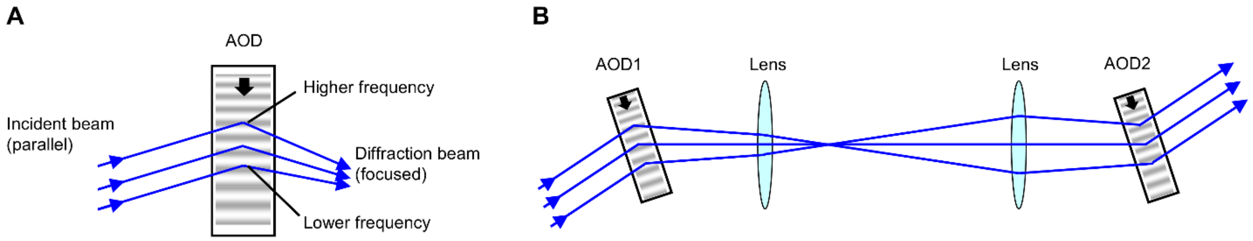

**Fig. S12. Beam scanning using acousto-optic deflectors.** (*A*) Illustration of beam focusing during high-speed scanning with a single acousto-optic deflector (AOD). (*B*) Dual-AOD configuration that cancels beam focusing at high scanning speeds.

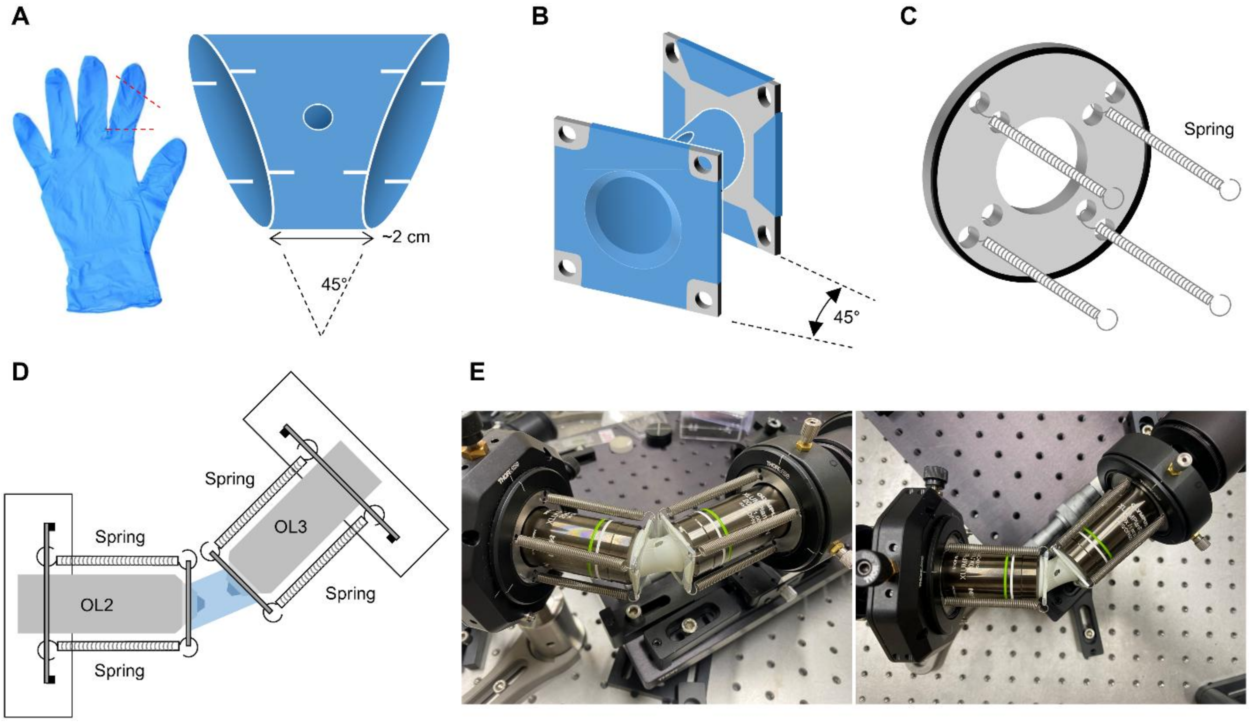


**Fig. S13. Water chamber.** (*A*) Main body of the chamber: A trapezoidal tube made from the finger portion of a rubber glove, with four notches on each end. (*B*) Main body of the chamber attached to metal plates. (*C*) Auxiliary jig for mounting the chamber onto the front end of the objective lenses. (*D*) Schematic of the chamber and auxiliary jig arrangement. The chamber is pressed against the objective lens to create a water-tight seal. (*E*) Photographs of an assembled chamber.

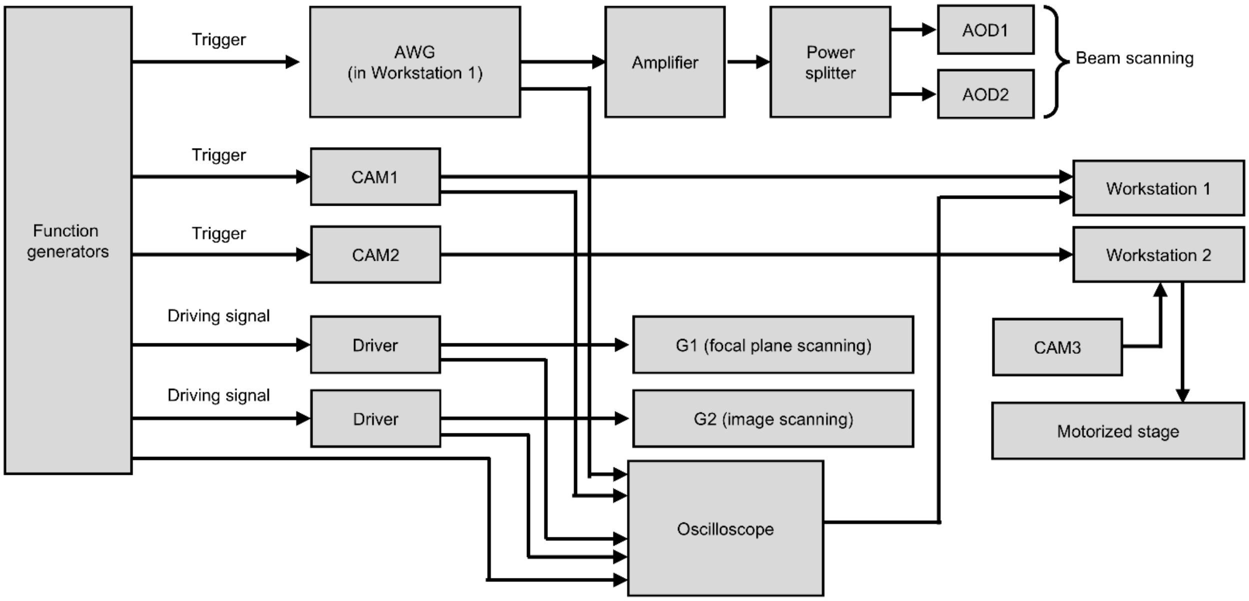


**Fig. S14. ISOP Microscope control system.** A series of function generators generate trigger and driving signals to synchronously control the acousto-optic deflectors (AOD1 and AOD2), galvanometric scanners (G1 and G2), and cameras (CAM1 and CAM2). Independently, a motorized stage and another camera (CAM3) are controlled to track target organisms. Monitoring signals are recorded by an oscilloscope, which play an essential role in the calibration process.

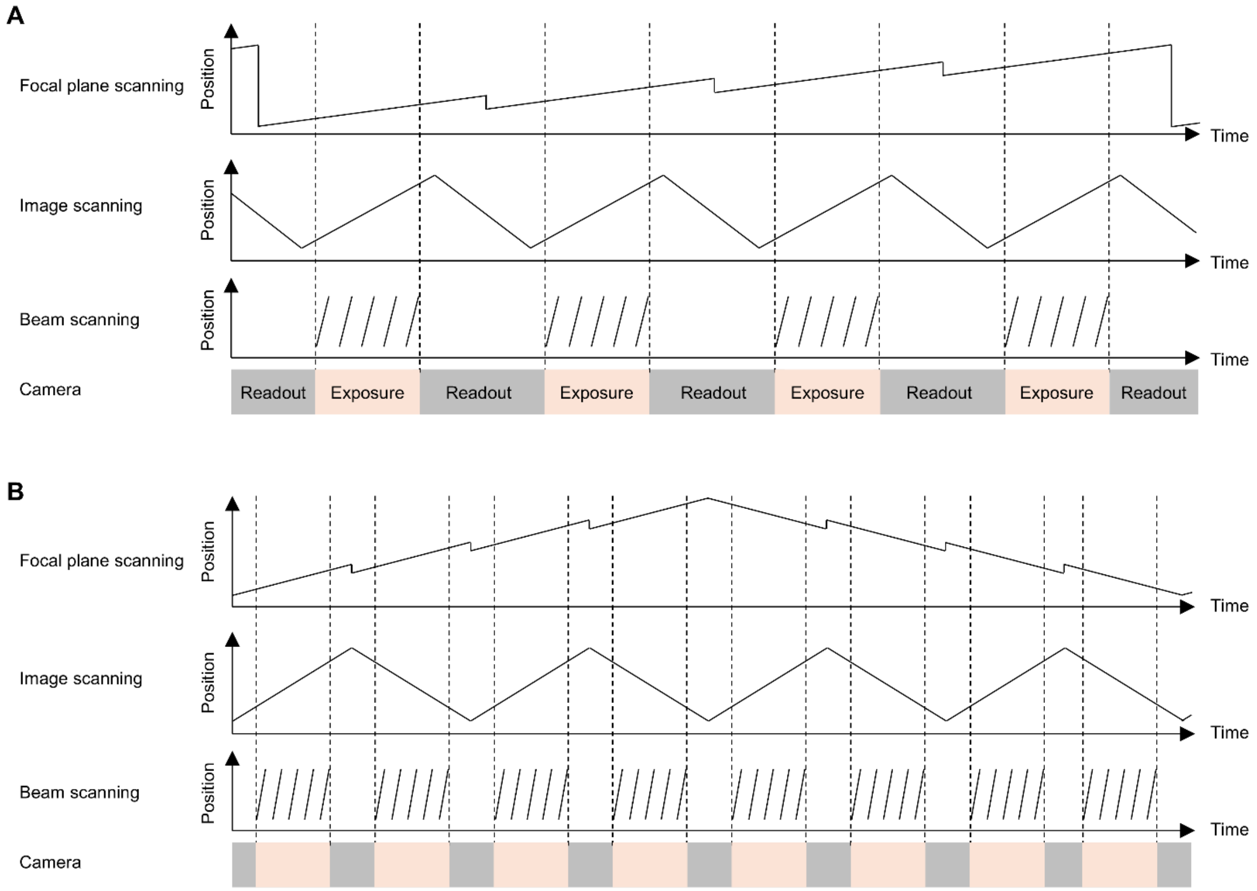


**Fig. S15. Typical scanning sequences and camera exposure timings.** (*A*) Scanning sequences and camera exposure timings for unidirectional image scanning (type I). (*B*) Scanning sequences and camera exposure timings for bidirectional image scanning (type II).

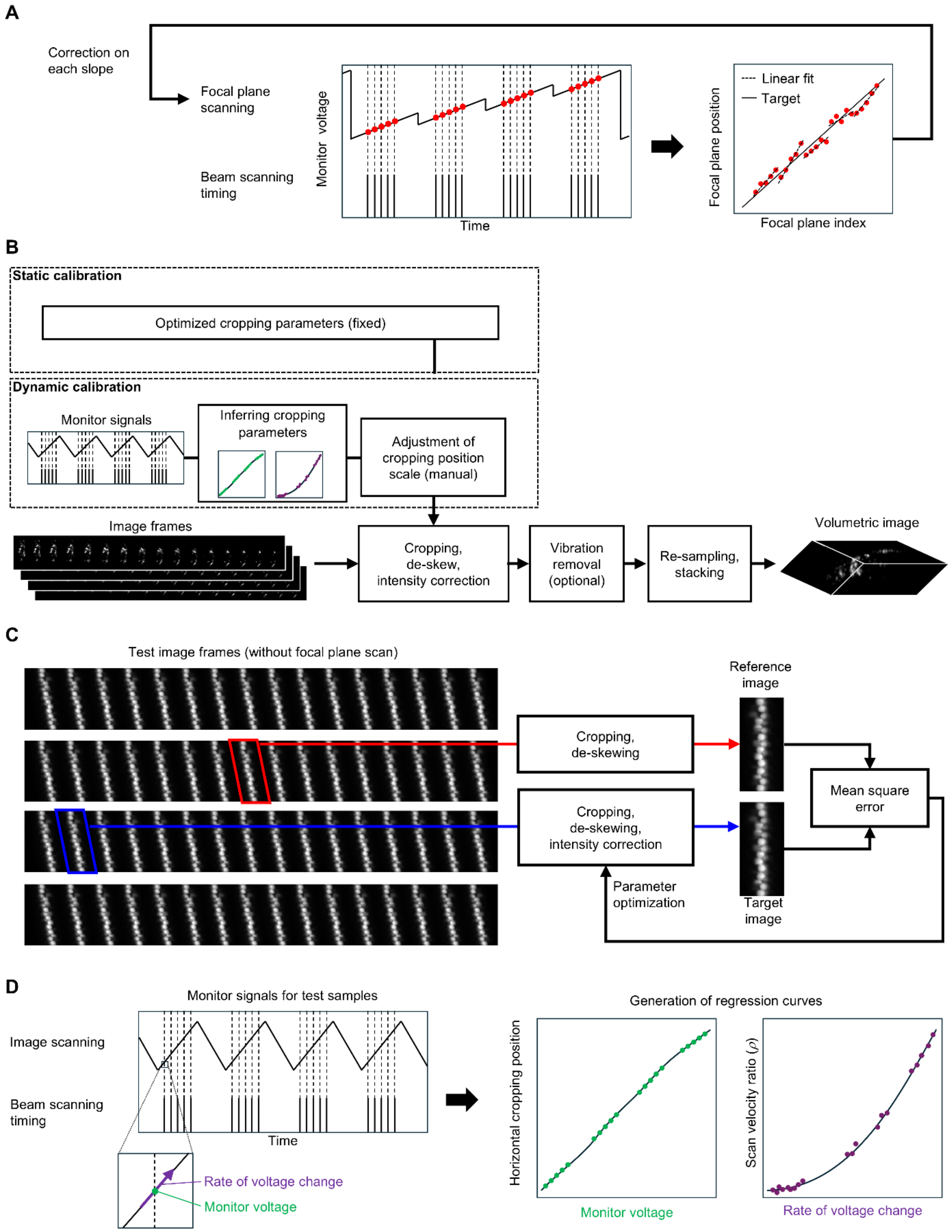


**Fig. S16. Calibration procedures for image acquisition and volumetric image reconstruction.** (*A*) Calibration procedure for focal plane scanning. (*B*) Volumetric image reconstruction from image frames. (*C*) Static calibration for extracting sectioned images. (*D*) Dynamic calibration for extracting sectioned images.

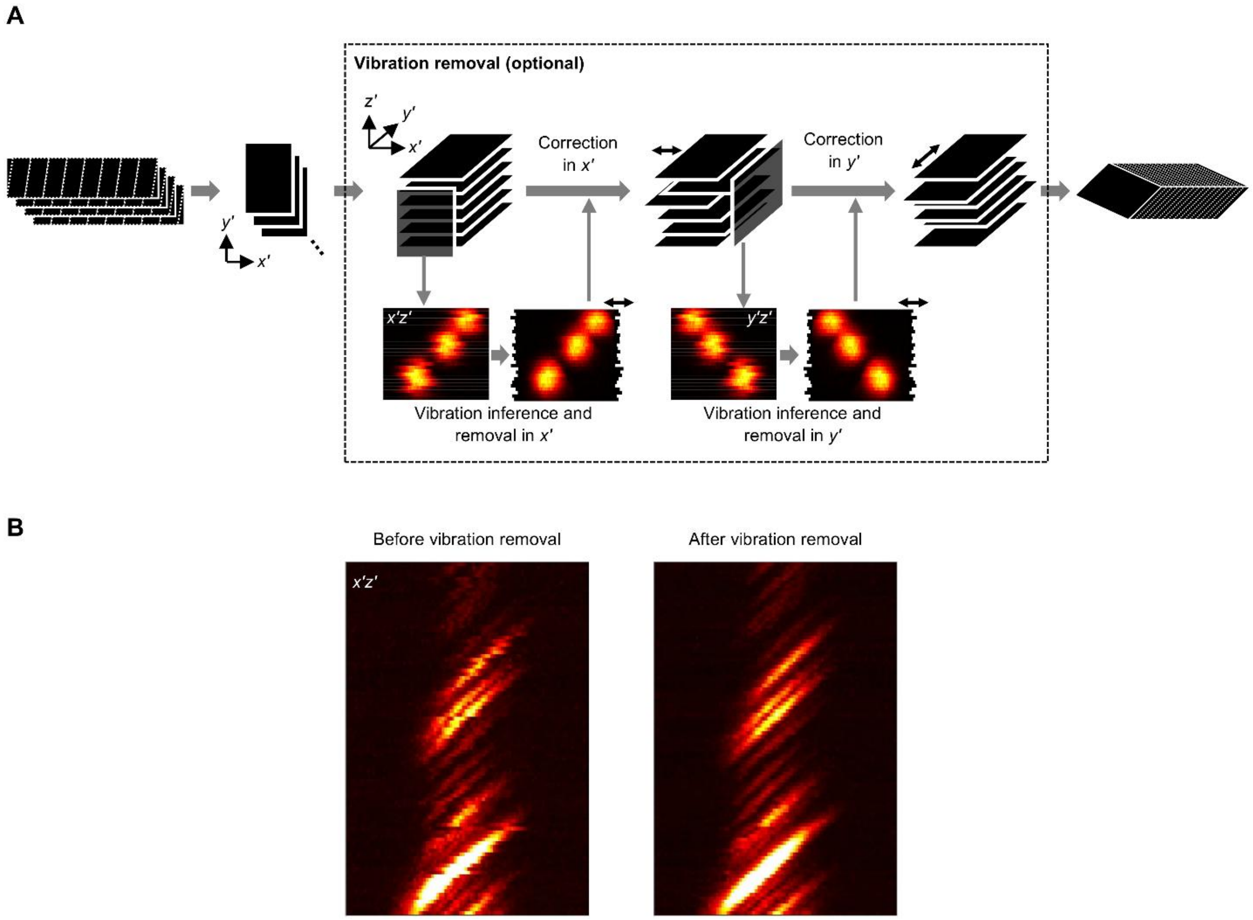


**Fig. S17. Vibration removal.** (*A*) Data processing workflow. (*B*) Maximum intensity projection images of 200-nm fluorescent beads in the *x′z′* plane before and after vibration removal.

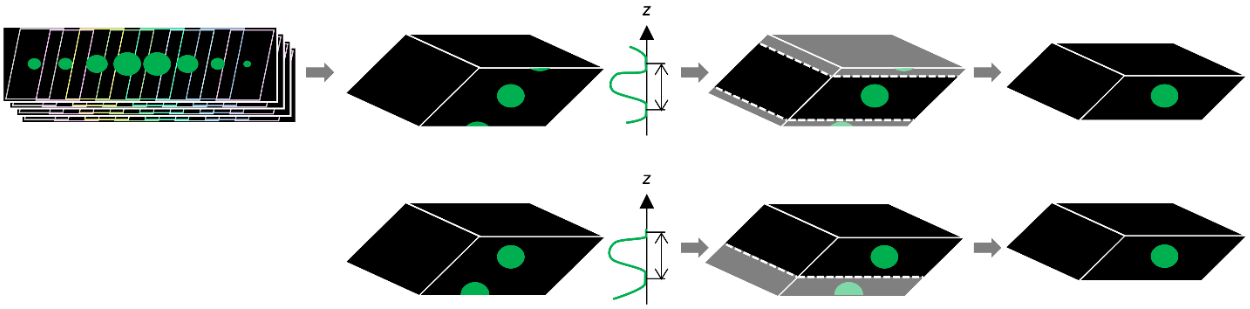


**Fig. S18. Virtual *z*-position correction.** A volumetric image is cropped from one with a broader depth range to exclude the folded regions of the object located at the top or bottom.

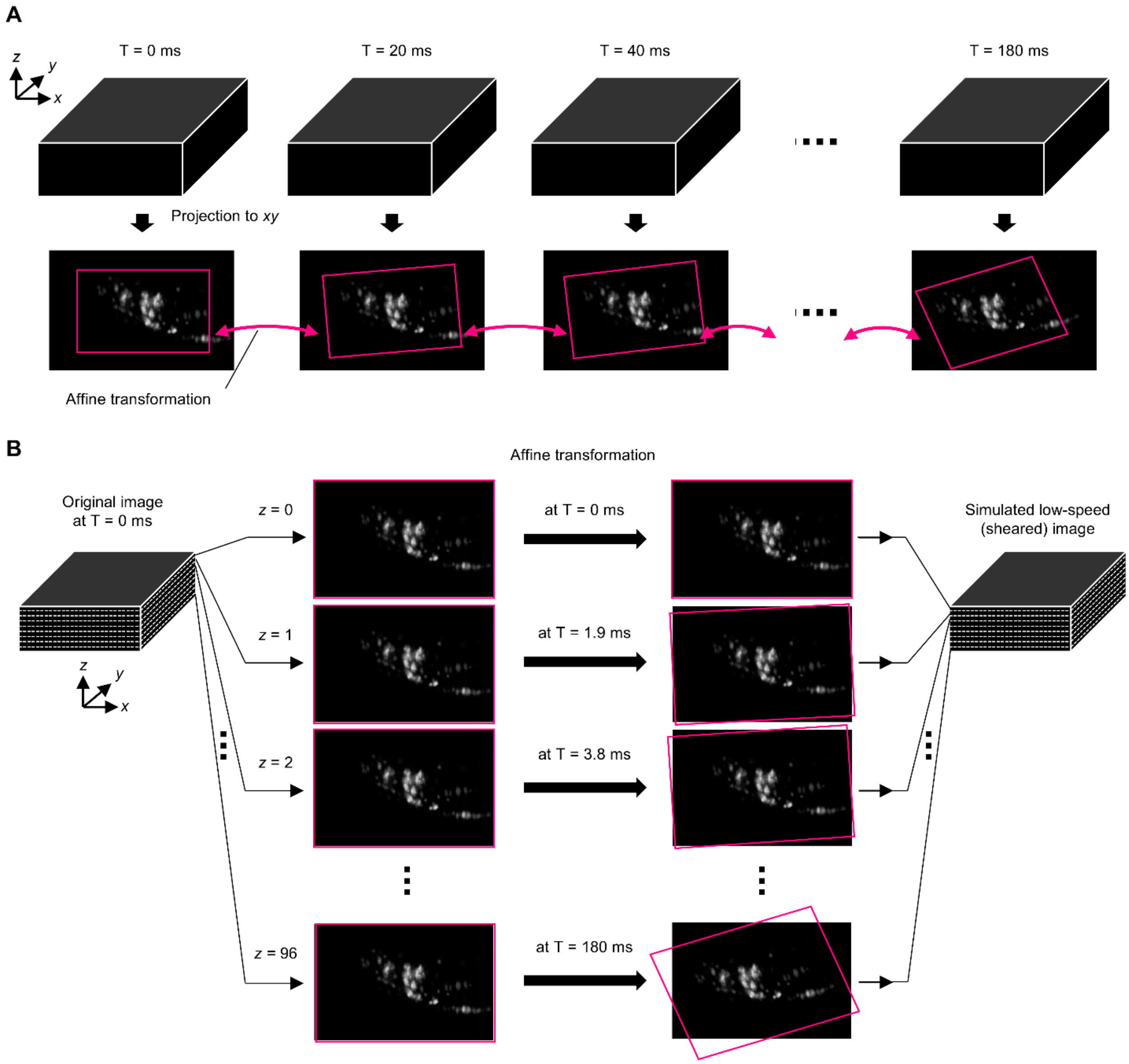


**Fig. S19. Generation of simulated low-speed imaging datasets.** (*A*) Prediction of worm posture changes at each time point using pattern matching between *xy*-projection images, represented by affine transformations. (*B*) Generation of simulated low-speed imaging dataset using the identified affine transformations.

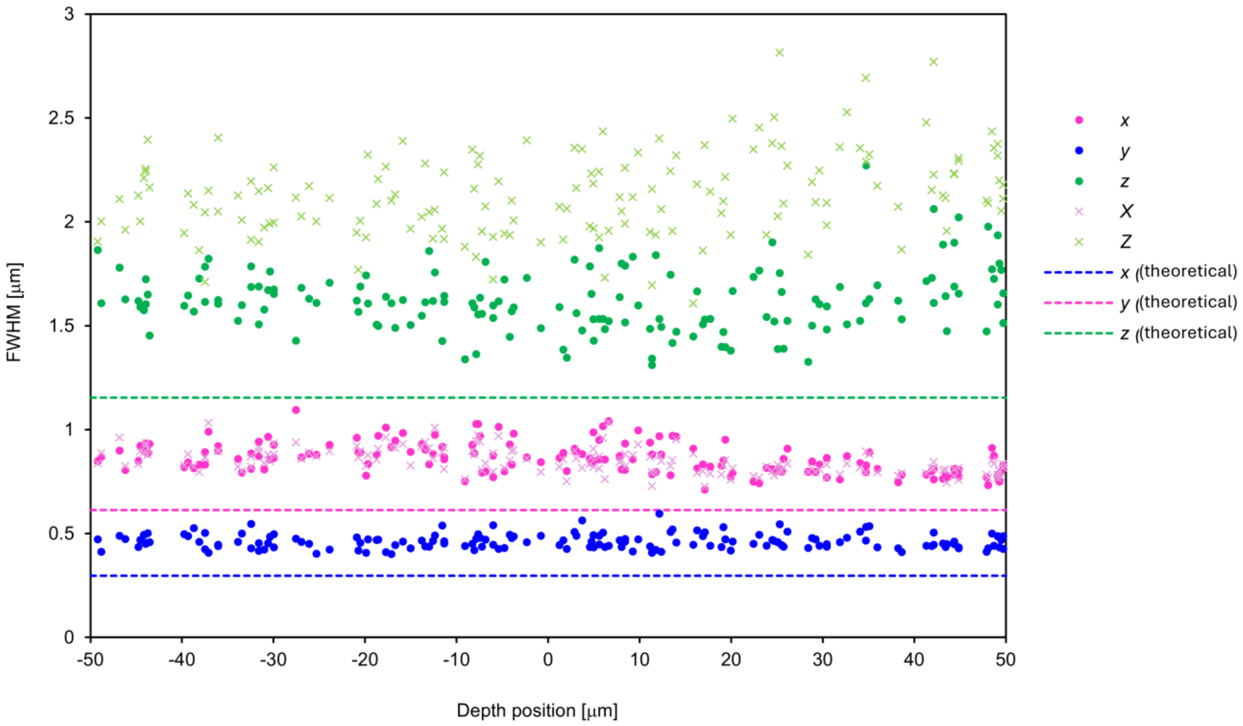


**Fig. S20. Point spread function (PSF) measurements within the ±50-μm depth range.** The vertical axis indicates the full width at half maximum of offset Gaussian functions fitted to line profiles obtained from images of 200-nm fluorescent beads after vibration removal. The plot was generated from 170 images in total. The *X* and *Z* axes represent orientations tilted by 23° relative to the *x* and *z* axes, respectively, corresponding to the minimum and maximum widths of theoretical PSFs in the *xz* plane. Theoretical values are based on PSFs numerically simulated using scalar diffraction theory with the Debye approximation.

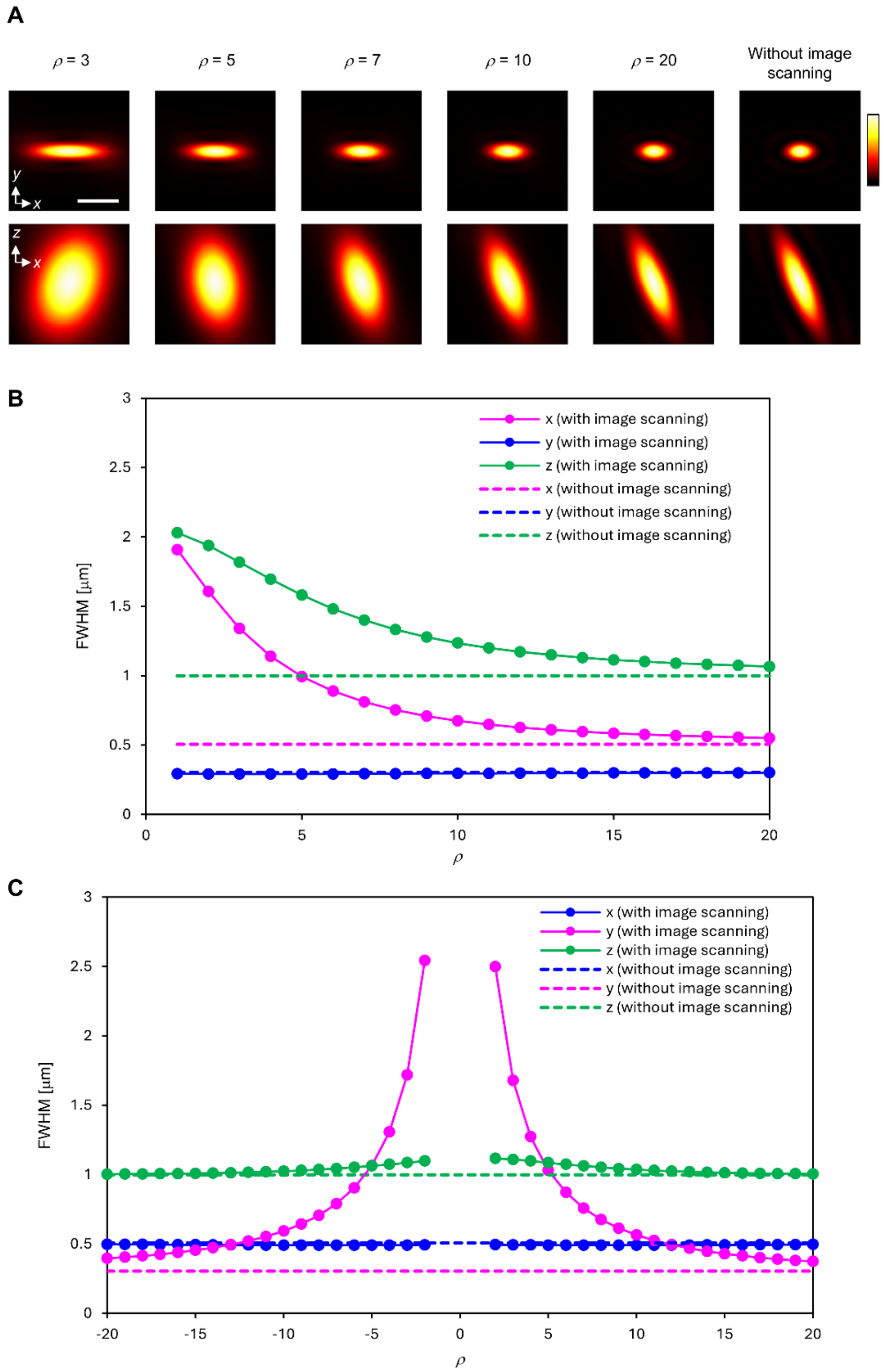


**Fig. S21. Characterization of theoretical point spread functions (PSFs).** (*A*) Intensity profiles of theoretical PSFs in the *xy* and *xz* planes at various scanning velocity ratios ($\rho$). (*B*) FWHM values of theoretical PSFs along the *x*, *y*, and *z* directions at various scanning velocity ratios ($\rho$) under a scanning configuration in which the beam scanning direction is perpendicular to the image scanning direction. Dashed lines indicate the theoretical values assuming no image scanning. (*C*) Same as (*B*), but with the beam scanning direction parallel or antiparallel to the image scanning direction.

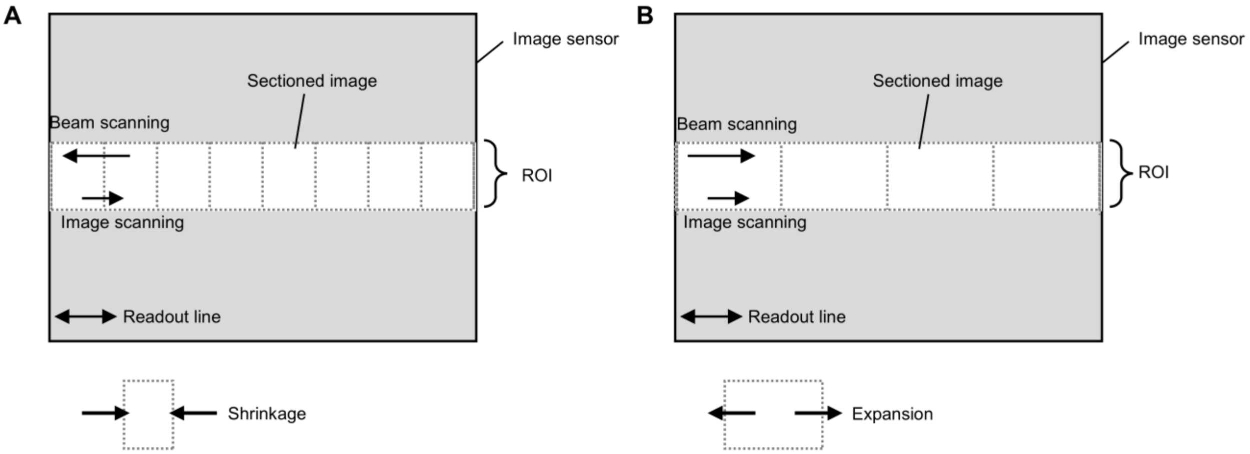


**Fig. S22. Alternative scanning configurations.** (*A*) Configuration in which the beam scanning direction is opposite to the image scanning direction. (*B*) Configuration in which the beam scanning direction is aligned with the image scanning direction.

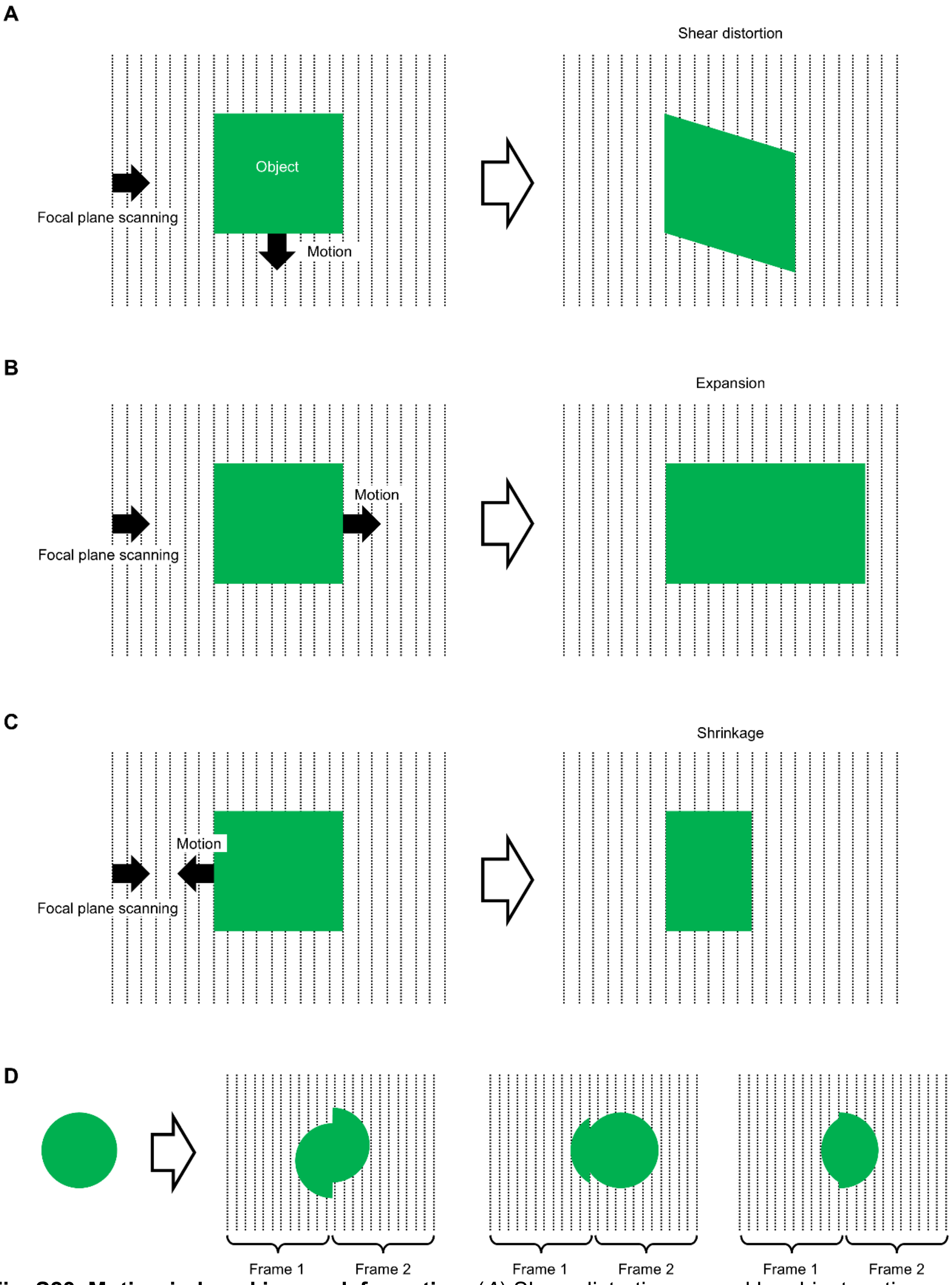


**Fig. S23. Motion-induced image deformation.** (*A*) Shear distortion caused by object motion orthogonal to the focal plane scanning direction. (*B*) Image expansion caused by object motion in the same direction as the focal plane scanning direction. (*C*) Image shrinkage caused by object motion opposite to the focal plane scanning direction. (*D*) Discontinuous image deformation at boundaries between local volumetric regions corresponding to consecutive image frames.

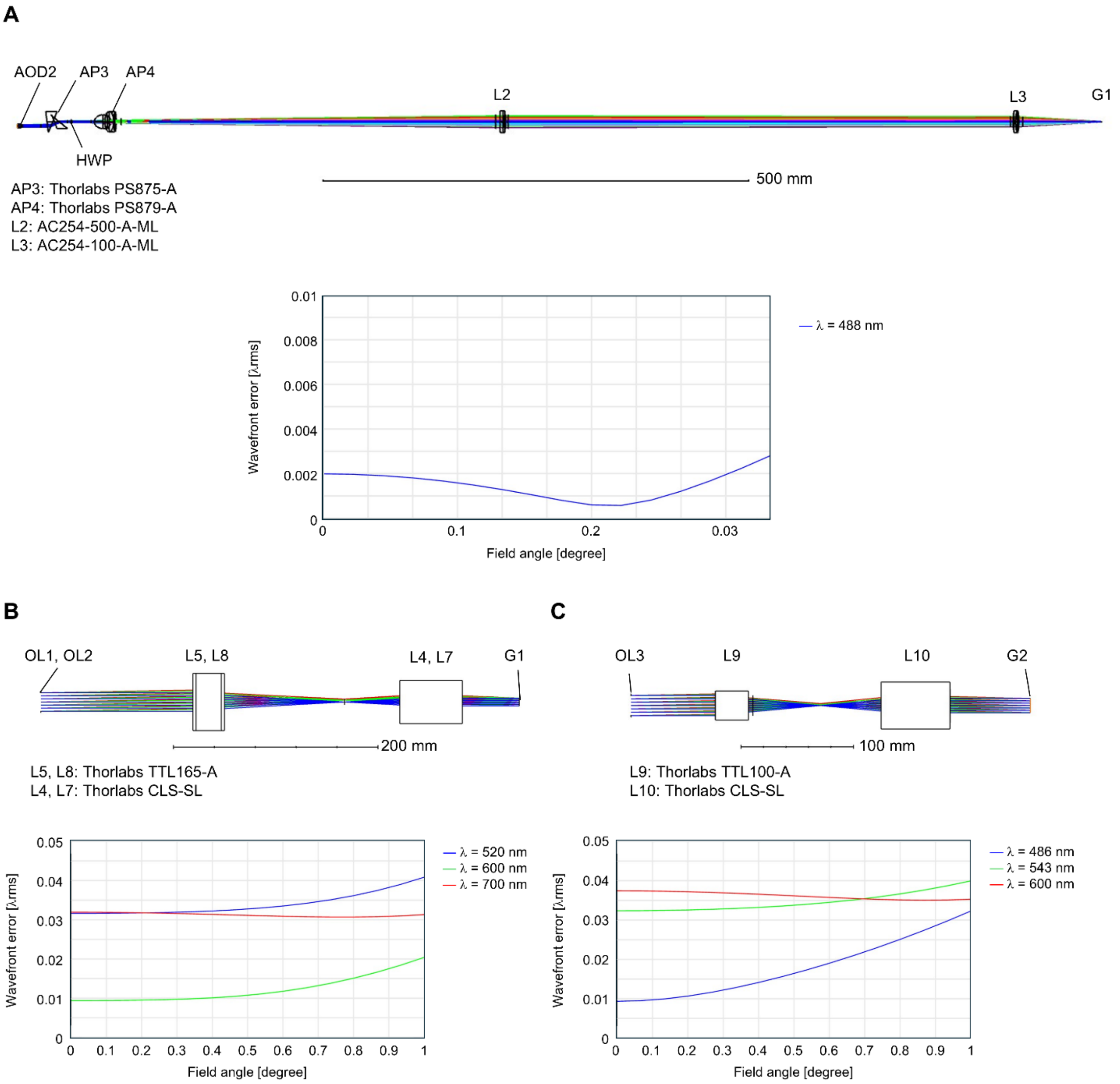


**Fig. S24. Optical design of relay optics in the ISOP microscope.** (*A*) Relay optics between AOD2 and G1 and their aberration characteristics. (*B*) Relay optics between G1 and OL1/OL2 and their aberration characteristics. (*C*) Relay optics between OL3 and G2 and their aberration characteristics.

| | *C. elegans* | *H. exemplaris* | *C. reinhardtii* | 200-nm fluorescent beads (for PSF evaluation) |
|---|---|---|---|---|
| Laser | Coherent Genesis CX488-2000 STM-SV | Coherent Genesis CX488-2000 STM-SV | Coherent Genesis CX488-2000 STM-SV | Coherent Genesis CX488-2000 STM-SV |
| AP1 | Thorlabs PS879-A | Thorlabs PS879-A | Thorlabs PS879-A | Thorlabs PS879-A |
| AP2 | Thorlabs PS875-A | Thorlabs PS875-A | Thorlabs PS875-A | Thorlabs PS875-A |
| AP3 | Thorlabs PS875-A | Thorlabs PS875-A | Thorlabs PS875-A | Thorlabs PS875-A |
| AP4 | Thorlabs PS875-A | Thorlabs PS875-A | Thorlabs PS875-A | Thorlabs PS875-A |
| AOD1 | Brimrose TED-150-100-488 | Brimrose TED-150-100-488 | Brimrose TED-150-100-488 | Brimrose TED-150-100-488 |
| AOD2 | Brimrose TED-150-100-488 | Brimrose TED-150-100-488 | Brimrose TED-150-100-488 | Brimrose TED-150-100-488 |
| L1 | Thorlabs AC254-100-A-ML | Thorlabs AC254-100-A-ML | Thorlabs AC254-100-A-ML | Thorlabs AC254-100-A-ML |
| L2 | Thorlabs AC254-100-A-ML | Thorlabs AC254-100-A-ML | Thorlabs AC254-100-A-ML | Thorlabs AC254-100-A-ML |
| L3 | Thorlabs AC254-500-A-ML | Thorlabs AC254-500-A-ML | Thorlabs AC254-500-A-ML | Thorlabs AC254-500-A-ML |
| L4 | Thorlabs AC254-100-A-ML | Thorlabs AC254-100-A-ML | Thorlabs AC254-100-A-ML | Thorlabs AC254-100-A-ML |
| L5 | Thorlabs CLS-SL | Thorlabs CLS-SL | Thorlabs CLS-SL | Thorlabs CLS-SL |
| L6 | Thorlabs TTL165-A | Thorlabs TTL165-A | Thorlabs TTL165-A | Thorlabs TTL165-A |
| L7 | Thorlabs CLS-SL | Thorlabs CLS-SL | Thorlabs CLS-SL | Thorlabs CLS-SL |
| L8 | Thorlabs TTL165-A | Thorlabs TTL165-A | Thorlabs TTL165-A | Thorlabs TTL165-A |
| L9 | Thorlabs TTL100-A | Thorlabs TTL100-A | Thorlabs TTL100-A | Thorlabs TTL100-A |
| L10 | Thorlabs CLS-SL | Thorlabs CLS-SL | Thorlabs CLS-SL | Thorlabs CLS-SL |
| L11 | Thorlabs TTL100-A | Thorlabs TTL100-A | Thorlabs TTL100-A | Thorlabs TTL100-A |
| L12 | Thorlabs TTL100-A | Thorlabs TTL100-A | Thorlabs TTL100-A | Thorlabs TTL200-A |
| L13 | Thorlabs AC254-125-B-ML | Thorlabs AC254-125-B-ML | Thorlabs AC254-125-B-ML | Thorlabs AC254-125-B-ML |
| L14 | Thorlabs AC254-200-B-ML | Thorlabs AC254-200-B-ML | Thorlabs AC254-200-B-ML | Thorlabs AC254-200-B-ML |
| OL1 | Olympus XLUMPLFLN 20XW | Olympus XLUMPLFLN 20XW | Olympus XLUMPLFLN 20XW | Olympus XLUMPLFLN 20XW |
| OL2 | Olympus XLUMPLFLN 20XW | Olympus XLUMPLFLN 20XW | Olympus XLUMPLFLN 20XW | Olympus XLUMPLFLN 20XW |
| OL3 | Olympus XLUMPLFLN 20XW | Olympus XLUMPLFLN 20XW | Olympus XLUMPLFLN 20XW | Olympus XLUMPLFLN 20XW |
| DM1 | Semrock Di01-R488/561 | Semrock Di01-R488/561 | Semrock Di01-R488/561 | Semrock Di01-R488/561 |
| DM2 | Semrock FF580-FDi01 | Semrock FF580-FDi01 | Semrock FF580-FDi01 | (Unused) |
| DM3 | Semrock FF699-FDi01-t3-25x36 | Semrock FF699-FDi01-t3-25x36 | Semrock FF699-FDi01-t3-25x36 | Semrock FF699-FDi01-t3-25x36 |
| Filters | Semrock NF03-488E-25 | Semrock NF03-488E-25 | Semrock NF03-488E-25 | Semrock NF03-488E-25 FF01-561/4-25 |
| G1 | Thorlabs GVS211/M | Thorlabs GVS211/M | Thorlabs GVS211/M | Thorlabs GVS211/M |
| G2 | Thorlabs GVS211/M | Thorlabs GVS211/M | ScannerMax Saturn 9B | Thorlabs GVS211/M |
| CAM1 | Hamamatsu ORCA Fusion C14440-20UP | Hamamatsu ORCA Fusion C14440-20UP | Hamamatsu ORCA Fusion C14440-20UP | Hamamatsu ORCA Fusion C14440-20UP |
| CAM2 | Hamamatsu ORCA Fusion C14440-20UP | Hamamatsu ORCA Fusion C14440-20UP | Hamamatsu ORCA Fusion C14440-20UP | (Unused) |
| CAM3 | Basler acA2040-120uc | Basler acA2040-120uc | (Unused) | (Unused) |
| Slit | Thorlabs VA100 | Thorlabs VA100 | Thorlabs VA100 | Thorlabs VA100 |
| Motorized stage | OptoSigma OSMS26-50(XY) | OptoSigma OSMS26-50(XY) | OptoSigma OSMS26-50(XY) | OptoSigma OSMS26-50(XY) |
| Function generators | Teledyne LeCroy Wavestation 2012, 2052, 2022; RIGOL DG400 | Teledyne LeCroy Wavestation 2012, 2052, 2022; RIGOL DG400 | Teledyne LeCroy Wavestation 2012, 2052, 2022; RIGOL DG400 | Teledyne LeCroy Wavestation 2012, 2052, 2022; RIGOL DG400 |
| Arbitrary waveform generator | Signatec PXDAC4800A-DP | Signatec PXDAC4800A-DP | Signatec PXDAC4800A-DP | Signatec PXDAC4800A-DP |
| USB oscilloscope | Pico Technology PicoScope 4824A | Pico Technology PicoScope 4824A | Pico Technology PicoScope 4824A | Pico Technology PicoScope 4824A |
| Amplifier for AOD | Mini-Circuits ZHL-1-2W+ | Mini-Circuits ZHL-1-2W+ | Mini-Circuits ZHL-1-2W+ | Mini-Circuits ZHL-1-2W+ |
| Power splitter | Mini-Circuits ZA2CS-251-20WS+ | Mini-Circuits ZA2CS-251-20WS+ | Mini-Circuits ZA2CS-251-20WS+ | Mini-Circuits ZA2CS-251-20WS |

**Table S1. List of optical and electronic components.**

| # | Sample | *C. elegans* | *H. exemplaris* | *C. reinhardtii* | 200-nm fluorescent beads (for PSF evaluation) |
|---|---|---|---|---|---|
| 1 | Image acquisition mode | type I | type I | type II | type I |
| 2 | Number of pixels in the image scanning direction* | 141 | 171 | 95 | 57 |
| 3 | Number of pixels in the direction perpendicular to image scanning | 300 | 708 | 124 | 60 |
| 4 | Number of sectioned images | 128 | 195 | 24 | 40 |
| 5 | Number of sectioned images per frame | 16 | 13 | 24 | 40 |
| 6 | Number of frames per volume | 8 | 15 | 1 | 1 |
| 7 | Global exposure time [ms] | 0.9 | 3 | 0.37 | 200 |
| 8 | $a_{G2}$ (see Materials and Methods for the definition) | 0.7 | 0.7 | 0.4 | 0.95 |
| 9 | Pixel size [μm] | 0.41 | 0.41 | 0.41 | 0.205 |
| 10 | Sectioned image spacing in the *x* direction [μm] | 1.3 | 1.5 | 3.05 | 0.4 |
| 11 | Ratio of beam scanning speed to image scanning speed ($\rho$) | 8.9 | 8.9 | 8.9 | 1.3–19.1 |
| 12 | Volume rate [Hz] | 50 | 10.2 | 1000 | 5 |
| 13 | FOV in the *x* direction [μm]** | 207 | 342 | 101 | 24 |
| 14 | FOV in the *y* direction [μm]** | 123 | 290 | 51 | 12 |
| 15 | FOV in the *z* direction [μm]** | 41 | 50 | 28 | 8 |
| 16 | Average laser power to the sample [mW] | 0.18 (Worm 4)<br>0.26 (Worm 1, 3)<br>0.32 (Worm 2) | 2.6 | 1.5 (for swimming cells shown in Fig. 4*A* and Movie S6)<br>4.8 (for flowing cells shown in Fig. 4*D* and Movie S7) | 0.8 (at scan velocity ratio of 8.9) |

*Slightly lower than the values shown in Fig. S2 because of finite beam scanning speed.
**Values calculated from 2–4, 9, and 10; actual FOVs may vary slightly due to fine adjustments in image reconstruction.

**Table S2. Imaging parameters for each dataset.**

| Time (Volume index) | Number of detected cells | | | | |
|---|---|---|---|---|---|
| | Manually detected | Detected by model_1 | Detected by model_3000 | Detected by model_6000 | Detected by model_9995 |
| 1 | 161 | 160 | 152 | 155 | 162 |
| 3000 | 153 | 138 | 152 | 135 | 147 |
| 6000 | 136 | 111 | 114 | 135 | 120 |
| 9995 | 136 | 116 | 122 | 123 | 120 |

**Table S3. Number of detected cells using 3D StarDist.** The numerical identifier in each model name (e.g., *model_1*) indicates the volumetric image used for training.

**Movie S1. Principles of image-scanning LSM.**
On the left, the process of image formation in image-scanning digital scanned light-sheet microscopy (DSLM) is illustrated, where both beam scanning and image scanning are employed. On the right, the process of image formation in image-scanning selective plane illumination microscopy (SPIM) is shown. In this case, stroboscopic illumination is used as an alternative to beam scanning.

**Movie S2. Whole-brain calcium imaging of freely behaving *C. elegans* (Worm 1).**
Maximum intensity projections in the *xy*, *xz*, and *zy* planes of dual-color fluorescence volumetric images acquired at 50 vps (left) and corresponding brightfield images (right) from the head of a freely behaving *C. elegans* (Worm 1). The dual-color images show neuronal nuclei co-expressing tdTomato (magenta) and GCaMP6f (green). Photobleaching correction was applied (see Methods). The videos are shown in real time. Scale bar, 20 µm.

**Movie S3. Whole-brain calcium imaging of freely behaving *C. elegans* (Worms 2-4).**
Maximum intensity projections in the *xy*, *xz*, and *zy* planes of dual-color fluorescence volumetric images acquired at 50 vps from the head of freely behaving *C. elegans* (Worms 2-4). The dual-color images show neuronal nuclei co-expressing tdTomato (magenta) and GCaMP6f (green). Photobleaching correction was applied (see Methods). The videos are shown in real time. Scale bar, 20 µm.

**Movie S4. Neuronal tracking in the head of a freely behaving *C. elegans* using ISOP microscopy and spinning-disk confocal microscopy.**
Maximum intensity projections (MIPs) in the *xy*, *xz*, and *zy* planes of tdTomato fluorescence volumetric images (left) and the corresponding cell tracking results (right) from the head of a freely behaving *C. elegans*, acquired using ISOP microscopy at 50 vps (top, Worm 1) and spinning-disk confocal microscopy at 3.2 vps (bottom). Photobleaching correction was applied (see Methods). The spinning-disk confocal images were resampled in the *z* direction with linear interpolation before MIP generation to better visualize motion-induced deformation. Cell tracking was semi-automatically performed using 3DeeCellTracker without manual correction in either dataset. Tracking entirely failed at 46 s in the dataset acquired by spinning-disk confocal microscopy.

**Movie S5. Volumetric imaging of muscle fibers in a tardigrade during locomotion.**
Maximum intensity projections in the *xy*, *xz*, and *zy* planes of dual-color fluorescence volumetric images acquired at 10 vps (left) and corresponding brightfield images (right) from a tardigrade. The dual-color images show muscle fibers co-expressing mCherry (red-biased magenta) and GCaMP6s (green). The videos are shown in real time. Scale bar, 50 µm.

**Movie S6. Volumetric imaging of swimming *C. reinhardtii* cells at 1,000 vps.**
Maximum intensity projections in the *xy*, *xz*, and *zy* planes (top) and stereographic views with the parallel viewing method (bottom) of dual-color fluorescence volumes of swimming *C. reinhardtii* cells captured using ISOP microscopy at 1,000 vps for 10 s. The nuclei (green) were stained with SYTO16 while chloroplasts (magenta) were visualized by autofluorescence. Scale bar, 10 µm.

**Movie S7. Volumetric imaging of flowing *C. reinhardtii* cells at 1,000 vps.**
Maximum intensity projections in the *xy*, *xz*, and *zy* planes (top) and stereographic views with the parallel viewing method (bottom) of dual-color fluorescence volumetric images of flowing *C. reinhardtii* cells captured using ISOP microscopy at 1,000 vps for 10 s. The nuclei (green) were stained with SYTO9 while chloroplasts (magenta) were visualized by autofluorescence. Scale bar, 10 µm.

**Movie S8. Real-time cell tracking in freely behaving *C. elegans*.**
Two neurons in tdTomato fluorescence volumetric images of Worm 2, acquired using ISOP microscopy at 50 vps, were tracked using a real-time cell tracking algorithm. The positions of the tracked cells are shown as squares overlaid on the top-down MIPs of the tdTomato fluorescence

volumetric images with photobleaching correction (see Methods). Cell tracking was maintained for 33.8 s before terminating due to tracking errors. The video is shown in real time. Scale bar, 20 µm.

**SI References**


1. C. J. R. Sheppard, Super-resolution in confocal imaging. *Optik (Stuttgart)* 80, 53–54 (1988).
2. A. G. York, P. Chandris, D. D. Nogare, J. Head, P. Wawrzusin, R. S. Fischer, A. Chitnis, H. Shroff, Instant super-resolution imaging in live cells and embryos via analog image processing. *Nat. Methods* 10, 1122–1126 (2013).
3. E. D. Diebold, B. W. Buckley, D. R. Gossett, B. Jalali, Digitally synthesized beat frequency multiplexing for sub-millisecond fluorescence microscopy. *Nat. Photonics* 7, 806–810 (2013).
4. H. Mikami, J. Harmon, H. Kobayashi, S. Hamad, Y. Wang, O. Iwata, K. Suzuki, T. Ito, Y. Aisaka, N. Kutsuna, K. Nagasawa, H. Watarai, Y. Ozeki, K. Goda, Ultrafast confocal fluorescence microscopy beyond the fluorescence lifetime limit. *Optica* 5, 117 (2018).
